# Soluble E-cadherin Drives Brain Metastasis in Inflammatory Breast Cancer

**DOI:** 10.1101/2025.06.18.660428

**Authors:** Xiaoding Hu, Yun Xiong, Emilly S Villodre, Huiming Zhang, Juhee Song, Natalie Fowlkes, Savitri Krishnamurthy, Marissa Rylander, Chandra Bartholomeusz, Debu Tripathy, The MDACC Inflammatory Breast Cancer Team, Wendy A Woodward, Junjie Chen, Bisrat G Debeb

**Affiliations:** Departments of Breast Medical Oncology, The University of Texas MD Anderson Cancer Center, Houston TX; Experimental Radiation Oncology, The University of Texas MD Anderson Cancer Center, Houston TX; Biostatistics The University of Texas MD Anderson Cancer Center, Houston TX; Veterinary Medicine and Surgery, The University of Texas MD Anderson Cancer Center, Houston TX; Pathology, The University of Texas MD Anderson Cancer Center, Houston TX; Breast Radiation Oncology, The University of Texas MD Anderson Cancer Center, Houston TX; MD Anderson Morgan Welch Inflammatory Breast Cancer Clinic and Research Program, The University of Texas MD Anderson Cancer Center, Houston TX; Department of Biomedical Engineering, The University of Texas at Austin, Austin, TX

**Author notes:** **Correspondence**: Bisrat G Debeb, DVM, PhD Associate Professor Department of Breast Medical Oncology Section of Translational Breast Cancer Research The Morgan Welch Inflammatory Breast Cancer Research Program and Clinic The University of Texas MD Anderson Cancer Center 6565 MD Anderson Blvd, Houston, TX 77030.

**Keywords:** Soluble E-cadherin, sEcad, Brain metastasis, Reactive astrocytes, XIAP, CXCL1/CXCL8-CXCR2 Axis, CXCR2 antagonist, Inflammatory breast cancer, IBC

## Abstract

The brain is a common site of relapse in inflammatory breast cancer (IBC), an E-cadherin positive, aggressive form of breast cancer. We found that elevated serum levels of soluble E-cadherin (sEcad), an 80-kDa fragment of E-cadherin, in patients with metastatic IBC correlated with poorer outcomes and increased rates of brain metastases. In our effort to understand the underlying mechanism, we discovered that sEcad binds to XIAP, an inhibitor of cell death, activating the pro-survival NF-kβ signaling in tumor cells. We also discovered that sEcad affects the tumor cell microenvironment by enhancing cancer cell adhesion to endothelial cells and inducing reactive astrocytosis in the brain. In addition, we found that sEcad-mediated reactive astrocytosis relies on the CXCL1/CXCL8-CXCR2 axis and treatment with a brain-permeable CXCR2 antagonist reduced brain metastatic burden and prolonged survival. These findings implicate sEcad in brain metastasis and provide new insights into potential therapeutic targets for IBC.

**Graphical Abstract:** 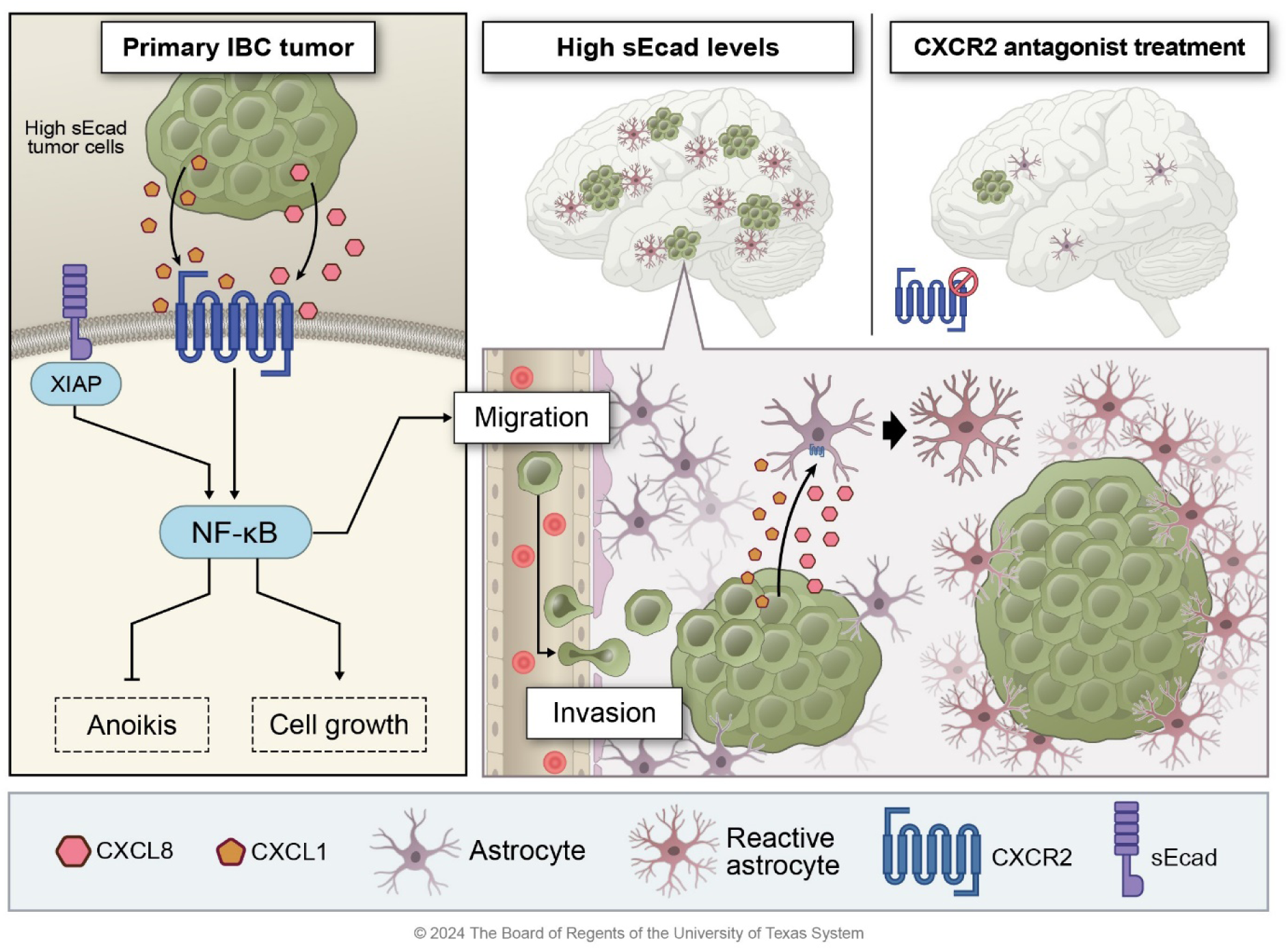

**Highlights:**

- High serum sEcad levels correlate clinically with poor survival outcomes and development of brain metastasis
- sEcad drives IBC brain metastasis growth in mouse models
- sEcad binds XIAP to activate NFkB and promote anoikis resistance and invasion of IBC cells
- sEcad activates reactive astrocytes and induces CXCR2 expression on tumor cells *in vitro* and *in vivo*
- CXCR2-IN-1, a brain-permeable CXCR2 antagonist, reduces metastasis and improves survival in IBC brain metastasis models

## INTRODUCTION

Brain metastasis is a common and challenging manifestation of solid malignancies such as lung, skin, and breast cancers, and is diagnosed in about 200,000 patients per year in the United States (ref). Brain metastasis occurs in 25%-50% of patients with advanced breast cancer, particularly those with the aggressive HER2-positive (HER2+) or triple-negative breast cancer (TNBC) subtypes (ref). Therapy options for brain metastases are limited and the current treatment portfolio is ineffective, and most patients die within months (ref), underscoring the critical need for new and effective treatments. However, the development of new therapies against brain metastasis is hampered by our limited understanding of the mechanisms that confer the competence of breast cancer cells during the numerous steps of the brain metastasis cascade. These include cancer cell migration and invasion from the site of the primary tumor into the bloodstream, survival within the high-stress circulatory environment, extravasation from blood capillaries into distant organs, and adaptation to the unique microenvironment of the brain.^1^

Brain metastasis is particularly common in patients with inflammatory breast cancer (IBC),^2,3^ a rare, highly aggressive form of breast cancer that in the United States accounts for 2%-4% of all breast cancer cases but contributes to up to 10% of breast cancer-related deaths.^4^ All patients with IBC present with lymph node involvement and one-third present with distant metastasis at diagnosis. HER2+ and TNBC subtypes are overrepresented in IBC,^5,6^ and more than 37% of patients with HER2+ IBC experience brain metastasis as the first site of relapse.^7^ IBC is currently treated with a multimodality approach consisting of systemic chemotherapy, surgery, and radiation therapy; however, the 5-year survival rates after this treatment range between 40 and 70%, depending on treatment location. A better understanding of the unique biology of IBC and identification of the molecular factors driving its aggressive and metastatic phenotype are urgently needed.

The overexpression of E-cadherin is a notable finding that distinguishes IBC from other breast cancers. E-cadherin expression has been reported as an indicator of low metastatic potential in many cancers; indeed, the loss of E-cadherin expression contributes to increased proliferation, invasion, and metastasis in some breast cancer models.^8,9^ However, recent studies have demonstrated a pro-metastatic role for E-cadherin.^10,11^ In IBC, E-cadherin is overexpressed in tumor cells, tumor emboli, metastases, and IBC cell lines.^12–17^ The presence of E-cadherin augments invasion and tumorigenesis in preclinical *in vitro* and *in vivo* IBC models^12,18–20^ and supports the formation of tumor emboli, a hallmark of IBC.^20,21^ These findings strongly support an oncogenic role for E-cadherin in IBC and possibly other tumors with high E-cadherin expression like ovarian cancer, glioma, invasive ductal carcinomas of the breast, gastric cancer and some subtypes of prostate cancer^22–31^.

E-cadherin is synthesized as a 120-kDa transmembrane glycoprotein; however, it can be cleaved off the ectodomain and released as a soluble form, designated soluble E-cadherin or sEcad. sEcad consists of an 80-kDa proteolytic fragment that is recognized as a biomarker of progression or recurrence in various cancers^32–35^. It has also shown tumor promoter functions through various mechanisms and activation of signaling pathways.^36–47^ For example, sEcad disrupts epithelial cell–cell adhesion, a major barrier against cancer cell mobility, thereby increasing the migration and invasion of cancer cells.^36,37^ It promotes disassembly of the E-cadherin–β-catenin adhesion complex and releases β-catenin to potentiate oncogenic Wnt signaling.^37–39^ sEcad also enhances cell survival by inhibiting apoptosis via functional interaction with membrane bound E-cadherin and activation of epidermal growth factor receptor (EGFR)– mediated PI3K/Akt/mTOR signaling.^40,47^ Further, sEcad increases the expression of metalloproteinases to promote tumor invasion into the stroma.^37,41^ Interestingly, sEcad may act as a soluble growth factor ligand that binds receptors such as HER1, HER2 and HER3 to activate EGFR signaling and promote aggressive tumor growth.^42–45^ sEcad can also be released within exosomes to promote tumor angiogenesis via activation of β-catenin and NFκB signaling.^46^ However, the roles of sEcad in breast cancer brain metastasis and IBC biology remain underexplored.

Here we report a direct correlation between elevated serum sEcad levels and increased development of brain metastasis and reduced survival in patients with metastatic IBC. We further identified sEcad as a key driver of brain metastasis in IBC mouse models. Mechanistically, we discovered that sEcad binds XIAP directly, activates NF-kβ signaling, and induces reactive astrocytosis through a targetable CXCL1/CXCL8-CXCR2 axis. Based on these findings, we propose a novel mechanism of brain metastasis in which sEcad functions as both a mediator of apoptotic resistance within tumor cells and a modulator of the brain microenvironment, thereby facilitating tumor cell survival, seeding, and outgrowth in the brain.

## RESULTS

### Serum sEcad levels correlate with increased development of brain metastasis and poor clinical outcomes in patients with IBC

In previous studies,^48^ we generated two sublines from a HER2+ metastatic IBC cell line, which we designated MDA-IBC3.1 (a weakly brain-metastatic subline) and MDA-IBC3.2 (a highly brain-metastatic subline). These sublines demonstrate distinct differences in their ability to metastasize to the brain after tail-vein injection into mice. Given that E-cadherin is consistently overexpressed in IBC tumor cells, tumor emboli, and metastases, and has been shown to have an oncogenic role in IBC,^13–17^ we examined the expression of E-cadherin in lysates from these two sublines. Interestingly, we observed remarkably higher expression of sEcad, but not full-length E-cadherin, in the highly brain-metastatic cells relative to the weakly brain-metastatic cells (Figure S1A). These findings were confirmed in the cell supernatants by enzyme-linked immunosorbent assay (ELISA) (Figure S1B), thus correlating sEcad to the propensity to form IBC brain metastasis in these sublines.

To examine the clinical relevance and overall significance of sEcad in patients with IBC, we measured serum levels of sEcad by ELISA in samples from 348 patients with IBC accrued to a prospective registry from the dedicated IBC clinic at The University of Texas MD Anderson Cancer Center (Table 1). We found that having higher serum sEcad levels correlated with poor overall survival (OS; *p*=0.04, Figure 1A), breast cancer specific survival (*p*=0.04, Figure 1B), earlier onset of metastasis (*p*=0.005, Figure 1C), and an increased risk of developing brain metastasis (*p*=0.04, Figure 1D). However, higher sEcad serum levels did not correlate with the incidence of lung metastasis (*p*=0.11, Figure S2A), suggesting sEcad’s potential role in organ site specific metastasis. Univariate analysis showed that sEcad levels, race, clinical disease stage, HR/HER2 status, lymphatic invasion, vascular invasion, and response to neoadjuvant chemotherapy were associated with overall survival and breast cancer–specific survival (Table 2). On multivariable analysis, high sEcad levels independently predicted poor OS (hazard ratio [HR]=2.07 [95% CI 1.19-3.60], *p*=0.009) along with clinical disease stage and receptor status (Figure 1E). These findings highlight the clinical significance of sEcad in patients with IBC and support its potential role in brain metastasis.

**Figure 1.**
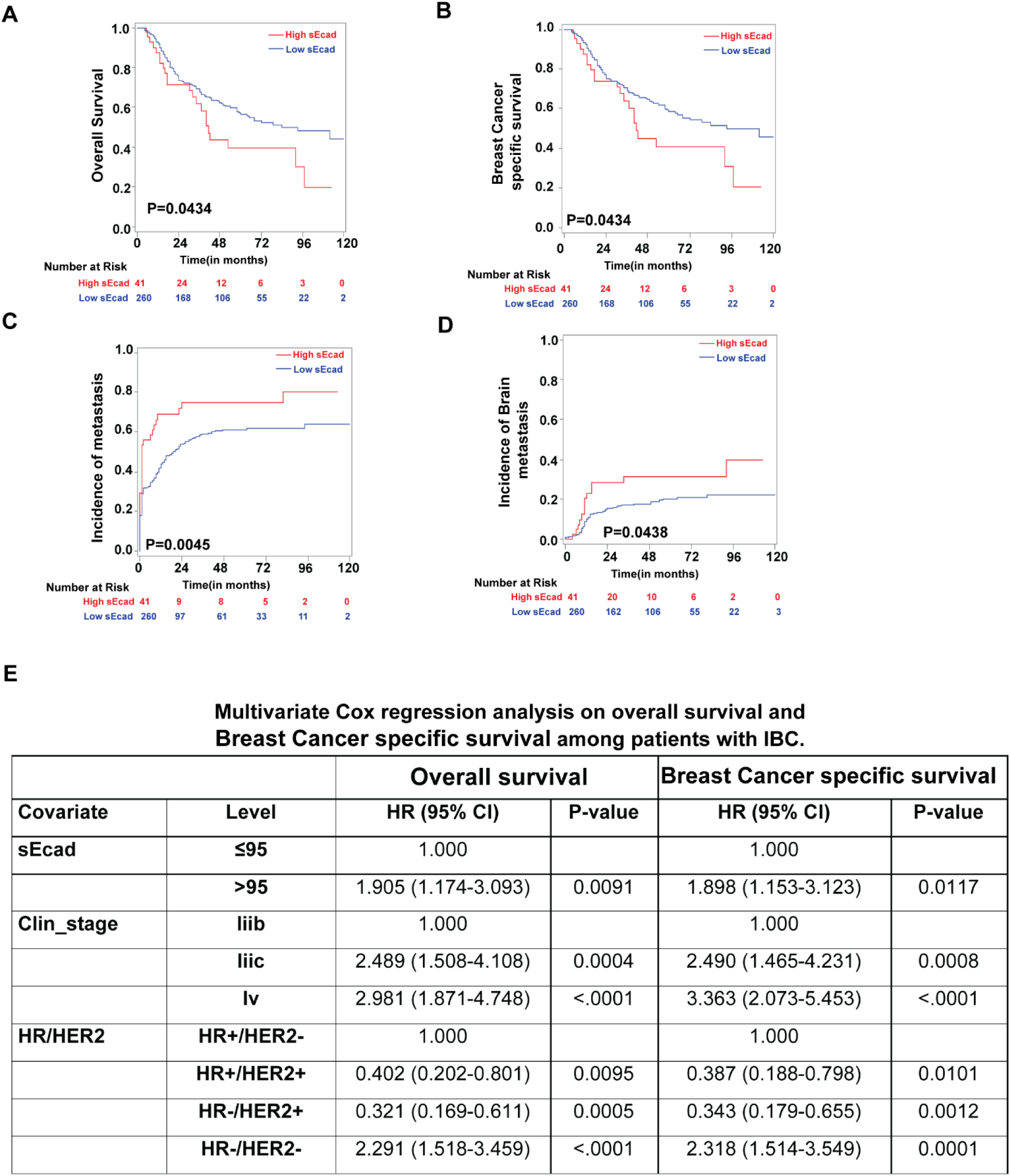
Higher serum sEcad levels clinically correlate with increased development of brain metastasis and poor survival outcomes in patients with inflammatory breast cancer (IBC) (A-D) Higher serum sEcad levels were associated with (A) reduced overall survival, (B) reduced breast cancer–specific survival, (C) increased risk of metastasis, and (D) increased risk of brain metastasis. (E) Multivariable analysis showing that higher serum sEcad level was independently correlated with poor survival outcome in 348 patients with IBC.

### sEcad promotes IBC cell invasion, migration, and resistance to anoikis

During metastatic progression, tumor cells show increased migratory and invasive abilities and resistance to cell death. Therefore, we examined whether sEcad affects these traits in IBC cells. We treated cells with sEcad recombinant protein in the presence or absence of DECMA1, a monoclonal neutralizing antibody against the ectodomain of E-cadherin, which has been shown to be inhibitory of sEcad.^49^ Treatment with sEcad recombinant protein (rsEcad) significantly increased anchorage-independent growth of IBC cells [rsEcad vs IgG control (MDA-IBC3, *p*=0.0009; SUM149, *p*=0.0005)], which was counteracted by DECMA1 [rsEcad+ DECMA1 vs rsEcad (MDA-IBC3, *p*=0.0046; SUM149, *p*=0.011)] (Figure 2A). sEcad also increased the size of soft agar colonies relative to the control group (MDA-IBC3, *p*<0.0001; SUM149, *p*=0.0061) (Figure 2B). These results indicate that sEcad-treated cells exhibit improved viability under non-adherent culture conditions. This anchorage-independent growth reflects the cancer cells’ ability to resist anoikis.^50–52^ Anoikis resistance, or the ability to escape apoptosis when detached from the extracellular matrix, is a crucial feature of cell survival during metastasis. Anoikis-resistant cells are known to form aggressive tumors that readily metastasize.^50,52^ Therefore, we examined the effects of sEcad on anoikis-mediated cell death in IBC cells. We found that treatment with recombinant sEcad significantly reduced cell death in IBC cells grown in poly-HEMA-treated low-attachment plates, compared with untreated controls [rsEcad vs IgG control (MDA-IBC3, *p*=0.0129; SUM149, *p*=0.0139)], an effect that was reversed by DECMA1 [sEcad+DECMA1 vs sEcad (MDA-IBC3, *p*=0.032; SUM149, *p*=0.0252)] (Figure 2C). These results suggest that sEcad enhances the resistance of breast cancer cells to anoikis, potentially enabling them to survive in circulation. We then examined the effect of sEcad on the migration and invasion of IBC cells by using the cell lines SUM149 and BCX010, which we used previously for similar studies as IBC3 has minimal baseline or induced migration capacity.^12^ Cells treated with recombinant sEcad exhibited increased migration (rsEcad vs IgG control [SUM149, *p*=0.0127; BCX010, *p*=0.0206]) (Figure 2D) and invasiveness (rsEcad vs IgG control [SUM149, *p*=0.0002; BCX010, *p*=0.0037]) (Figure 2E). Collectively, these findings underscore the possible significance of sEcad in driving the metastatic progression of IBC tumors.

**Figure 2.**
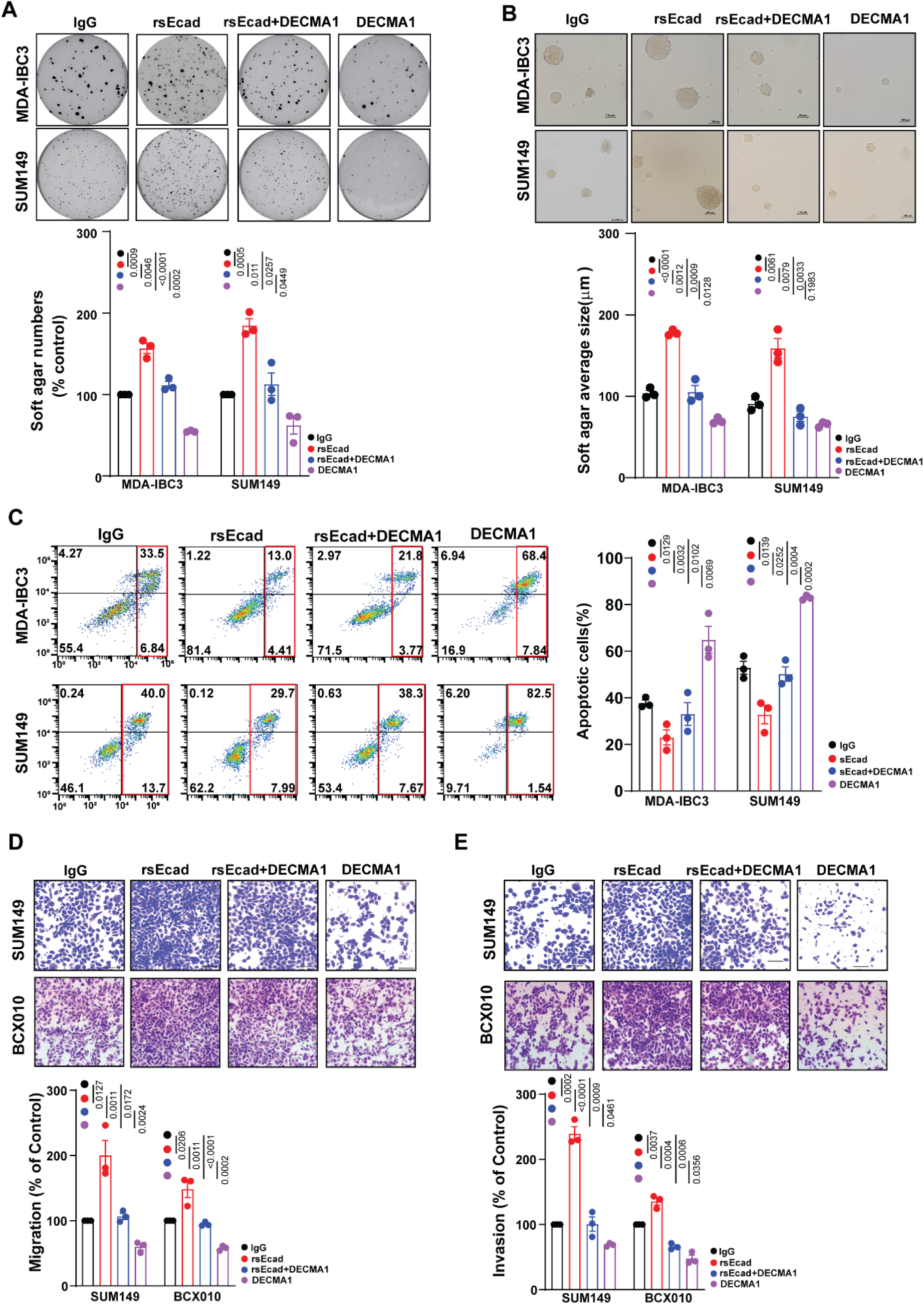
sEcad promotes IBC cell invasion, migration and resistance to anoikis. Inflammatory breast cancer (IBC) cells were treated with sEcad-recombinant protein (rsEcad) (20μg/mL) in the presence of absence of DECMA1 neutralizing antibody (20μg/mL). Data in (A-E) are mean ± SEM of three biological replicates. (A-B) rsEcad protein increased the (A) number and (B) size of anchorage-independent/soft agar-growing colonies of MDA-IBC3 and SUM149 cells, which was attenuated by treatment with DECMA1. (C) rsEcad protein led to increased anoikis resistance in MDA-IBC3 and SUM149 cells, which was reversed by DECMA1. (D-E) rsEcad protein enhanced the (D) migration and (E) invasion of SUM149 and BCX010 IBC cells, which were counteracted by the neutralizing antibody DECMA1.

As above, published studies of sEcad in cancer have primarily used recombinant proteins; however, a more stable and convenient cell-based system is essential to investigate sEcad’s role in tumor growth and metastasis and elucidate its underlying mechanisms. To address this need, we cloned the 80-kDa sEcad fragment based on the full-length E-cadherin structure (Figure S3A, left) into the Lenti-Blast vector and established stable Flag-tagged sEcad-overexpressing IBC cell lines [ER–/HER2+ (MDA-IBC3; SUM190) and ER–/HER2– (SUM149; BCX010)], which were validated by immunoblotting (Figure S3A, right). Using these newly established sEcad-overexpressing stable cell lines, we examined sEcad’s role in invasion, migration, anchorage-independent growth, and anoikis resistance. The results mirrored those observed with the sEcad recombinant protein, with overexpression of sEcad leading to a significant increase in the number and size of soft agar colonies in all four IBC cell lines (Figure S3B, S3C). Moreover, IBC cells overexpressing sEcad exhibited increased resistance to anoikis (Figure S3D) as well as increased migration (Figure S3E) and invasion (Figure S3F) compared with control groups.

In addition to the effects of sEcad on invasion and anoikis resistance, we examined whether sEcad enhances tumor cell competence for other steps in the brain metastasis cascade, including endothelial cell adhesion, angiogenesis, and trans-endothelial migration. Our findings show that sEcad-expressing IBC cells significantly increased tumor cell adhesion to endothelial cells (Figure S4A) and promoted angiogenesis (Figure S4B). Similar results were observed with sEcad recombinant protein, and these effects were neutralized by DECMA1 (Figure S4C). Moreover, in our *in vitro* blood-brain barrier (BBB) trans-endothelial migration model, sEcad-overexpressing SUM149 cells, demonstrated greater penetration across the BBB than the control (Figure S4D), suggesting that sEcad promotes the ability of tumor cells to invade the central nervous system tissue. Collectively, these findings validate our newly generated stable sEcad-overexpressing cell lines as a reliable model for studying the role of sEcad in tumor growth and metastasis. The results replicate our findings from the recombinant protein studies while providing a more stable, reproducible, and efficient system for mechanistic studies and animal experiments.

### sEcad is a driver of brain metastasis in IBC models

To determine the role of sEcad in brain metastatic progression, we used two IBC cell line models with a high propensity for brain metastasis (MDA-IBC3 and SUM149)^53^ with and without sEcad-Flag stable expression (Figure 3A). Mice injected via tail-vein with sEcad-overexpressing MDA-IBC3 cells had a higher incidence of brain metastasis compared with controls (MDA-IBC3-sEcad vs MDA-IBC3-Con: 100% vs 50%, *p*=0.032; Figure 3B). sEcad overexpressing IBC cells also formed larger, grossly visible brain metastases and increased both metastasis number (MDA-IBC3: *p*=0.0009, Figure 3C) and size (MDA-IBC3: *p*=0.0042, Figure 3D). Further, sEcad reduced the overall survival of mice (Figure 3E, *p*=0.0002) and shortened brain metastasis-specific survival time (MDA-IBC3: *p*=0.04, Figure 3F). In a second model, mice injected intracardially with sEcad-overexpressing SUM149 cells also had a higher incidence of brain metastasis (100% vs 60%; Figure 3B, lower panel), increased number of metastases (*p*=0.0023, Figure 3G), increased size of metastases (*p*=0.0015, Figure 3H), and reduced overall survival (*p*=0.0002, Figure 3I) and brain metastasis–free survival (*p*=0.001, Figure3J) compared with controls. Representative hematoxylin-and-eosin and immunohistochemically stained images of Flag and Ki67 (Figure 3K, 3L) confirmed sEcad overexpression and increased proliferation in the brain metastasis lesions in both the MDA-IBC3 and SUM149 models.

**Figure 3.**
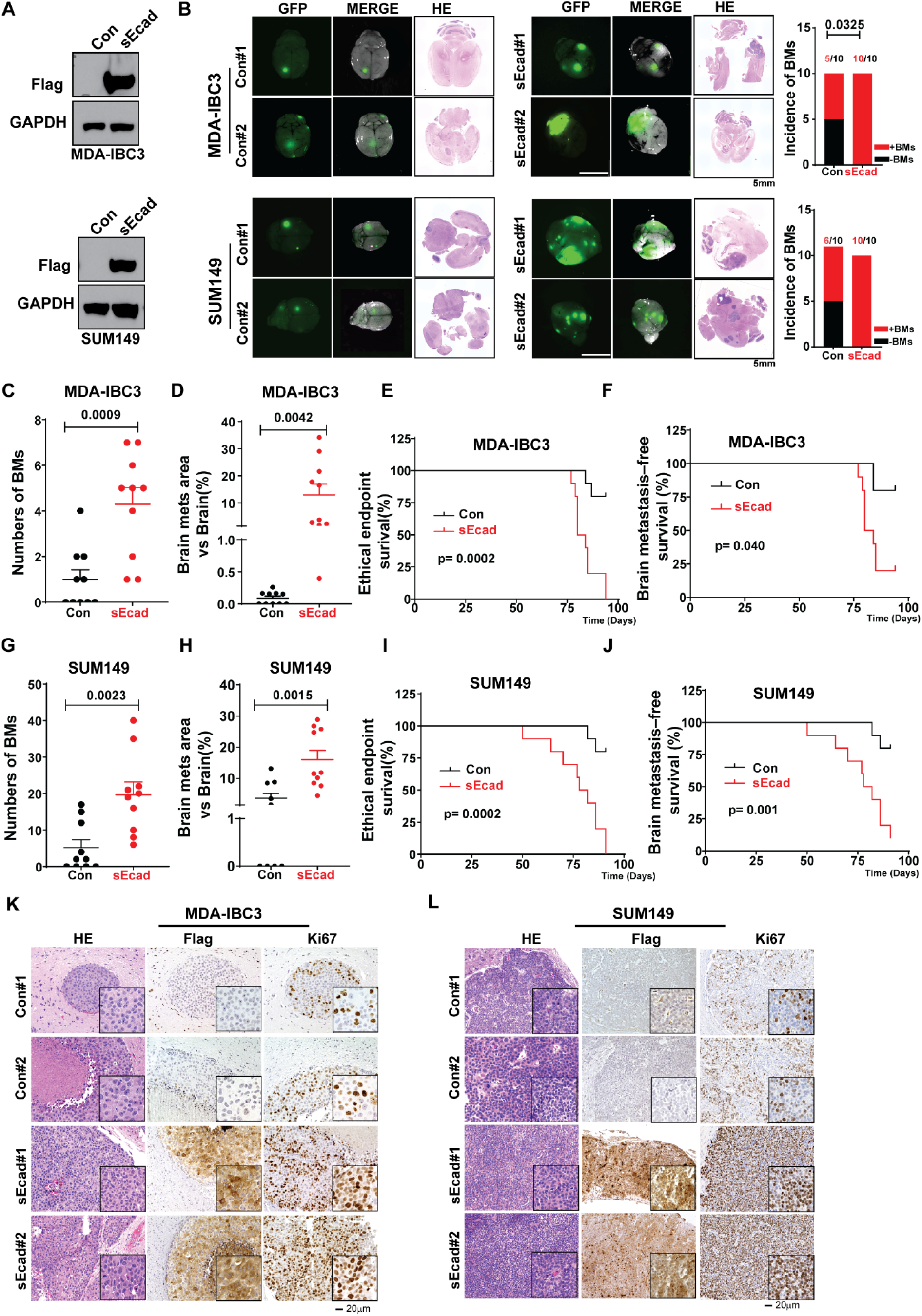
sEcad is a driver of brain metastasis in IBC models. (A) Generation of sEcad-FLAG overexpressing MDA-IBC3 and SUM149 stable IBC cell lines. Total cell lysates were analyzed by western blotting with anti-FLAG and anti-GAPDH antibodies. (B-L) For brain metastasis studies, HER2+ MDA-IBC3 cells (1 × 10^6^) were injected via tail vein, and the triple-negative IBC cell line SUM149 (2 × 10^5^) were injected via intracardiac injection into SCID/Beige mice. (B). (left) Representative images of brain metastases from sEcad-expressing and control MDA-IBC3 and SUM149 cells; (right) Incidence of brain metastases in sEcad-overexpressing MDA-IBC3 and SUM149 cells compared with controls. (C, G) Numbers of brain metastases in sEcad-overexpressing group compared with controls. (D, H) Size of brain metastasis lesions in sEcad-overexpressing group compared with controls. (E, I) Overall survival rates in mice injected with sEcad-overexpressing or control IBC cells. (F, J) Brain metastasis−free survival rates in mice injected with sEcad-overexpressing or control IBC cells. (K, L) Hematoxylin-and-eosin and immunohistochemical staining of brain metastasis lesions from MDA-IBC3 and SUM149 control cells or sEcad-overexpressing cells. Flag indicates sEcad overexpression, whereas Ki67 indicates proliferation in brain metastasis lesions from the two mouse models of IBC brain metastasis.

### sEcad interacts with the BIR2 domain of XIAP and activates NF-κB signaling

To elucidate the mechanism by which sEcad inhibits apoptosis and promotes anoikis resistance in IBC cells, we first analyzed the expression of pro- and anti-apoptotic proteins by using a human apoptosis proteomics array. This analysis revealed a marked upregulation of several anti-apoptotic proteins in sEcad-overexpressing MDA-IBC3 and SUM149 cells, with X-linked inhibitor of apoptosis protein (XIAP) showing the most significant increase (Figure 4A). To further investigate the molecular mechanisms underlying sEcad’s role in anoikis resistance, we sought to identify novel interacting protein partners. For this purpose, we cloned the 80-kDa sEcad fragment into a mammalian expression vector containing a 2×Strep-tag II affinity tag, generating stable SFB-tagged sEcad-expressing HEK293T cells. These cells were used for TAP-MS (tandem affinity-purification–mass spectrometry) and BioID (proximity-dependent biotinylation identification) –based proteomic profiling of the sEcad interactome. BioID has been used previously for detecting insoluble proteins or weak and/or transient interactions.^54,55^ Our mass spectrometry analysis identified several novel high-confidence sEcad-interacting proteins, with XIAP emerging as a top hit (false discovery rate [FDR] <0.01, SAINT score >0.8, and enrichment >50-fold; Figure 4B). BioID analysis further confirmed XIAP’s strong enrichment in the sEcad-high group (Figure 4C). The detection of XIAP by both TAP-MS and BioID reinforced its potential role in sEcad-mediated inhibition of apoptosis in IBC cells.

**Figure 4.**
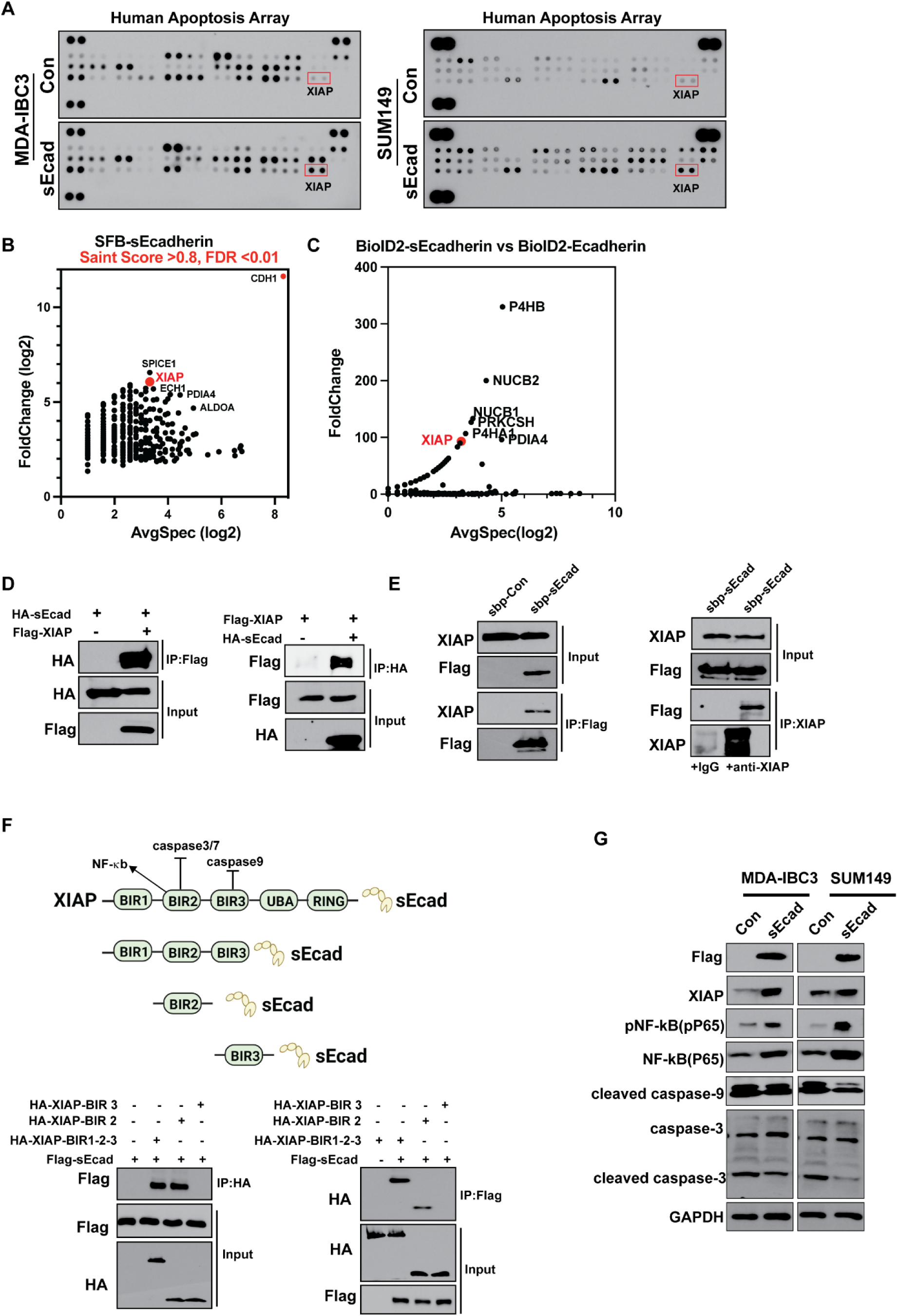
sEcad interacts with the BIR2 domain of XIAP and activates the NFκB pathway. (A) Human apoptosis array showed upregulation of XIAP in IBC cells that overexpress sEcad compared with controls. (B) Mass spectrometry and (C) BioID assays identified XIAP as a novel interaction partner of sEcad. (D) Exogenous immunoprecipitation assay confirmed the interaction between sEcad and XIAP. (E) Endogenous immunoprecipitation assay confirmed the interaction between sEcad and XIAP. (F) Domain mapping demonstrates that sEcad interacted with the BIR2 domain of XIAP. (G) Immunoblotting shows that sEcad enhanced XIAP expression, activated NFκB signaling, and inhibited the cleavage of caspase-3 in IBC cells.

To confirm the interaction between sEcad and XIAP under physiological conditions, we used reciprocal immunoprecipitation (IP) of the exogenous and endogenous proteins. Exogenously expressed HA-tagged sEcad interacted with Flag-XIAP *in vitro*, and vice versa (Figure 4D). Endogenous XIAP also associated with sEcad, as demonstrated by co-immunoprecipitation (co-IP) and reverse co-IP with an anti-XIAP antibody (Figure 4E). Through immunofluorescence experiments, we showed that sEcad is present in the cytoplasm of cancer cells, using both the cloned sEcad fragment (Fig. S5A) and the recombinant sEcad protein (Fig. S5B). These findings support the notion that sEcad is internalized by tumor cells, enabling its interaction with cytoplasmic targets such as XIAP. To map the XIAP domains involved in the sEcad-XIAP interaction, we used truncation mutants of XIAP [(HA-XIAP(BIR2), HA-XIAP(BIR3), and HA-XIAP(BIR1-BIR2-BIR3) obtained from Addgene]. As shown in Figure 4F, XIAP constructs containing the BIR2 domain, but not the BIR3 domain, pulled down sEcad, identifying the BIR2 domain as the region crucial for its interaction with sEcad. Others have shown that the BIR2 domain of XIAP primarily exerts its anti-apoptotic effects by regulating the NFκB pathway^56^ and caspase 3/7. ^57,58^ Consistent with this, our findings show that sEcad upregulates XIAP expression, activates NFκB signaling, and inhibits cleaved caspase 3 (Figure 4G), strongly supporting the role of sEcad in apoptosis resistance via the XIAP-NFκB pathway.

### sEcad triggers astrocyte reactivity

The anoikis resistance observed in rsEcad-treated and sEcad-overexpressing IBC cells (Figure 2 and Figure S3) could be associated with increased overall metastasis. However, our *in vivo* and patient data (Figures 1 and 3) suggest that sEcad preferentially drives brain metastases. This led us to hypothesize that in addition to cell autonomous induction of anoikis resistance, sEcad may have paracrine effects on the brain microenvironment inducing a supportive niche for metastatic breast cancer growth. Increasing evidence suggests that reactive astrocytes, marked by high levels of glial fibrillary acidic protein (GFAP), foster brain metastasis growth and progression via modulation of the blood-tumor barrier, promotion of an inflammatory and immunosuppressive brain microenvironment, enrichment of cancer stem cells, and increasing proliferation and invasion of metastatic cancer cells.^59–64^ To determine whether sEcad induces reactive astrocytosis, we treated an immortalized normal human astrocyte (NHA) cell line with recombinant sEcad (20 µg/mL for 36 h) or an IgG control. Immunofluorescence and immunoblotting analysis showed that sEcad significantly increased GFAP protein expression (Figure S6A and S6B), indicating direct induction of reactive astrocytes by sECAD *in vitro*. *In vivo* tail-vein injection of sEcad-overexpressing MDA-IBC3 cells led to a similar significant increase in GFAP-positive astrocytes within brain metastatic lesions relative to controls (Figure S6C). Similarly, intracardiac injection of SUM149-sEcad cells resulted in increased reactive astrocyte induction in brain tissue lesions compared to controls (Figure S6D). These findings were further validated by immunohistochemical staining with an anti-GFAP antibody (Figure S6E and S6F). These results suggest that tumor secreted sEcad acts on the microenvironment to promote a metastatic niche in the brain via induction of reactive astrocytes.

### sEcad-high cells induce CXCR2 activation via CXCL1/CXCL8

To elucidate the mechanisms by which sEcad can lead to modification of local microenvironments (in the brain or elsewhere), we performed a cytokine array analysis of secreted factors in conditioned medium collected from sEcad-expressing and control SUM149 and MDA-IBC3 cells. We found a significant increase in pro-inflammatory cytokines (CXCL1, CXCL8, and CXCL10) and secretory proteins (DKK1) in conditioned medium from sEcad-expressing cells (Figure 5A, Figure S7A). These findings were validated by ELISA (Figure 5B, Figure S7B).

**Figure 5.**
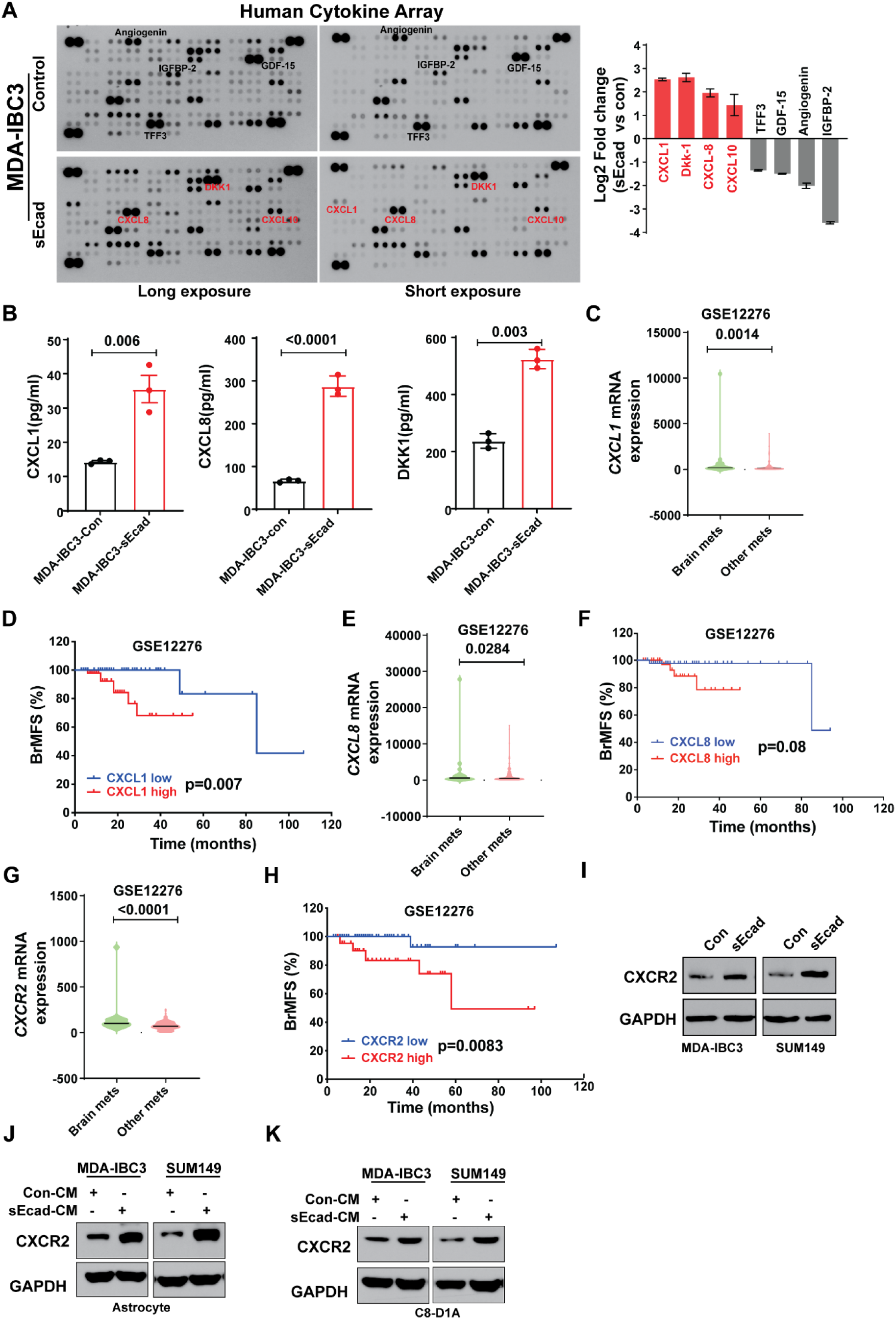
sEcad-overexpressing cells induce astrocyte and CXCR2 activation via CXCL1/CXCL8. (A, B) Cytokine array analysis of secreted factors showed increased levels of the pro-inflammatory cytokines CXCL1/CXCL8 in conditioned medium from sEcad-overexpressing MDA-IBC3 cells, as validated by ELISA. (C) CXCL1 was expressed at higher levels in patients with brain metastasis than in patients with metastasis to other anatomic sites. (D) Patients with high CXCL1 expression had reduced brain metastasis–free survival (BrMFs, (GSE12276); CXCL1-low indicates the bottom (25th) tertile, and CXCL1-high the top (75th) tertile. (E) CXCL8 was expressed at higher levels in patients with brain metastasis relative to patients with metastasis to other anatomic sites. (F) Patients with high CXCL8 expression had reduced brain metastasis*–*free survival (BrMFs, (GSE12276); CXCL8-low indicates the bottom (25th) tertile, and CXCL8-high the top (75th) tertile. (G) CXCR2 was expressed at higher levels in patients with brain metastasis than in patients with metastasis to other anatomic sites. (H) Patients with high CXCR2 expression had reduced brain metastasis–free survival (BrMFs, (GSE12276); CXCL2-low indicates the bottom (25th) tertile, and CXCL2-high the top (75th) tertile. (I) CXCR2 expression was induced in sEcad-overexpressing MDA-IBC3 and SUM149 cells relative to control cells. (J, K) Induction of CXCR2 expression in human astrocytes (J) and mouse astrocytes (K) treated with 30% condition medium from sEcad-overexpressing cells. GAPDH served as a loading control.

To prioritize these based on clinical relevance, we analyzed gene expression data obtained from patients with metastatic breast cancer (GEO dataset GSE12276).^65^ We found that the expression of CXCL1 and CXCL8, but not DKK1, is significantly higher in primary tumors that developed brain metastases compared with primary tumors that developed other metastases (Figure 5C, 5E; Figure S8A). Moreover, having high CXCL1 or CXCL8 expression, but not DKK1 expression, is correlated with reduced brain metastasis–free survival (Figure 5D, 5F, Figure S8B), suggesting that CXCL1 and CXCL8 may be related to sEcad’s promotion of brain metastases. Notably, expression of the CXCR2 receptor, a common receptor for both CXCL1 and CXCL8, was also significantly higher in patients with brain metastases and correlated with reduced brain metastasis–free survival (Figure 5G, 5H). These findings were intriguing given that CXCR2 is expressed in cancer cells, astrocytes, and microglia,^66^ suggesting that the CXCL1/CXCL8-CXCR2 axis may have a pivotal role in the brain-metastatic preference and progression of sEcad-expressing IBC cells. Consistent with these findings, we found increased expression of CXCR2 in cancer cells (Figure 5I) and in human and mouse astrocytes (Figure 5J, 5K) treated with conditioned medium from sEcad-overexpressing IBC cells compared with those treated with conditioned medium from control cells. CXCR2 was expressed at consistently higher levels in brain metastasis lesions and surrounding brain parenchyma from sEcad-high MDA-IBC3 cells relative to those from control cells (Figure 6A), with similar data obtained in brain tissues from SUM149 sEcad-high cells (Figure 6B). These observations implicate the CXCL1/CXCL8-CXCR2 axis in promoting sECAD-mediated reactive astrocytosis and brain metastasis.

**Figure 6.**
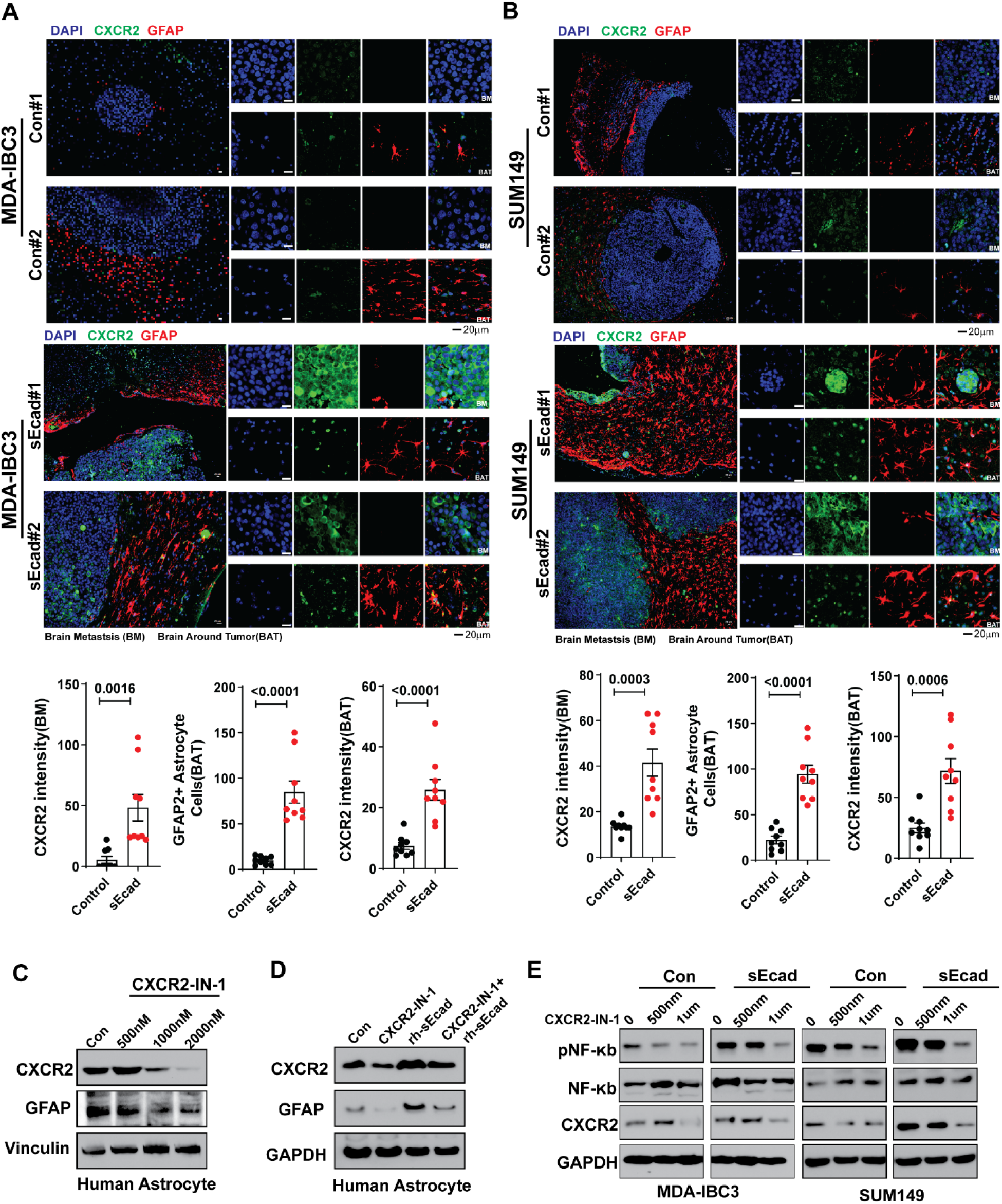
sEcad induces reactive astrocytes and CXCR2. (A, B) Immunofluorescence staining showed that metastatic lesions from sEcad-expressing tumors had significantly higher GFAP+ reactive astrocytes and CXCR2 expression than did the Control groups. Quantitation of CXCR2, GFAP-positive astrocytes. Data are mean ± SEM, n = 3 mice and 3 fields per mouse: t test. (C) Western blotting shows that the CXCR2 inhibitor CXCR2-IN-1 decreased CXCR2 and GFAP levels in human astrocytes; Vinculin served as a loading control. (D) Inhibition of CXCR2 and GFAP by CXCR2-IN-1 could be rescued by sEcad-recombinant protein (rsEcad). GAPDH served as a loading control. (E) sEcad increased the expression of CXCR2 and phosphorylated NFκB (pP65) in IBC cells, both of which were inhibited by CXCR2-IN-1. GAPDH served as a loading control.

### CXCR2 inhibitor significantly reduces brain metastasis and prolongs survival

To examine if sEcad promotes reactive astrocytes and brain metastases via a CXCL1/CXCL8-CXCR2 related mechanism, we inhibited this axis using CXCR2-IN-1, a brain-penetrant inhibitor of CXCR2,^67^ that may have potential as a pharmacologic tool for studying brain metastasis. Treatment of normal human astrocytes with various doses of CXCR2-IN-1 resulted in reduced expression of CXCR2 and GFAP, as confirmed by western blotting and immunofluorescence (Figure 6C, Figure S9A). Further, CXCR2-IN-1 reduced GFAP+ reactive astrocytes induced by sEcad recombinant protein (Figure 6D, Figure S9B). Similar results were obtained with another CXCR2 antagonist, SB225002, which effectively inhibited CXCR2 and GFAP expression (Figure S9C, S9D). In sEcad-high IBC cells, both CXCR2-IN-1 and SB225002 suppressed the phosphorylation of NFκB (p65) (Figure 6E, Figure S9E). In addition, colony-formation in sEcad-overexpressing MDA-IBC3 cells (Figure S10A) and SUM149 cells (Figure S10B) treated with CXCR-IN-1 was significantly reduced compared with vehicle controls. Similarly, enhanced migration (Figure S10C) and invasion (Figure S10D) of SUM149 sEcad-overexpressing cells were blocked by CXCR-IN-1. These findings highlight CXCR2-IN-1 as a pharmacologic inhibitor of sEcad mediated reactive astrocytosis and tumor cell proliferation, migration, and invasion.

To investigate if CXCR2-IN-1 can similarly inhibit growth and progression of sEcad-expressing IBC using our brain metastasis models, we implemented two treatment strategies: an ’early treatment’ approach targeting microscopic metastases and a ’late treatment’ approach targeting macroscopic metastases, as outlined in Figure S11A. In the ’early treatment’ group, CXCR2-IN-1 significantly reduced brain metastases in both MDA-IBC3-sEcad and SUM149-sEcad models (Figures 7A, 7F). Quantitative analysis revealed significant decreases in the number of brain metastases per mouse (Figure 7B, 7G) and the incidence of brain metastasis (Figure 7C, 7H). CXCR2-IN-1 treatment also led to improved overall survival (Figure 7D, 7I) and brain metastasis-free survival (Figure 7E, 7J) in both models. Representative hematoxylin and eosin stains showing results of this treatment strategy are shown in Figure S11D.

**Figure 7.**
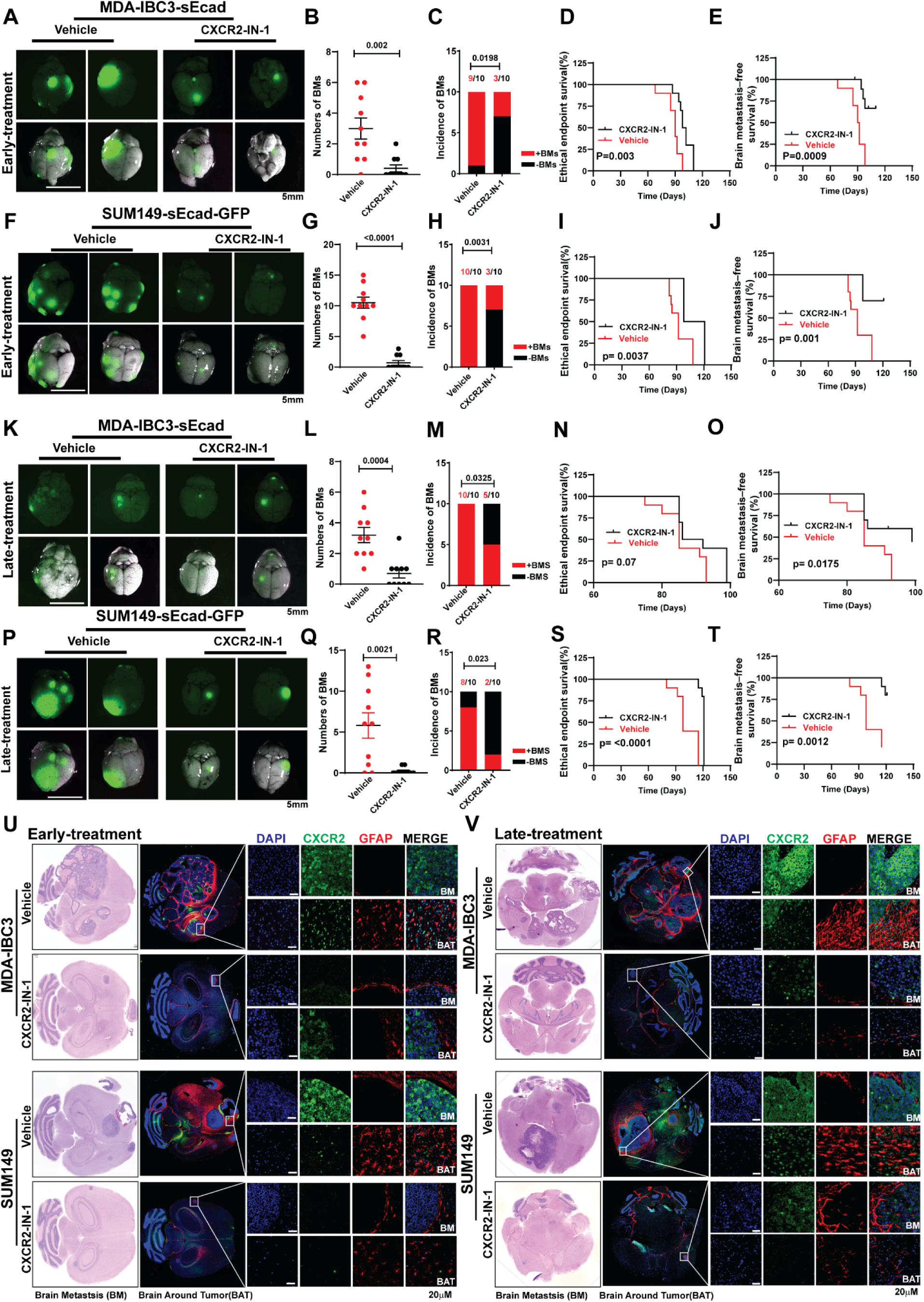
CXCR2 inhibitor reduces brain metastasis and prolongs survival. The schema for the ’Early’ and ’Late treatment’ approaches is outlined in Figure S11A. (A-J) Early Treatment group. CXCR2-IN-1 reduced brain metastases in both MDA-IBC3-sEcad and SUM149-sEcad models. (A, F) Representative images showed reduced brain metastasis in CXCR2-IN-1-treated MDA-IBC3-sEcad (A) and SUM149-sEcad (F) brain metastasis. CXCR2-IN-1 reduced the number of brain metastasis lesions (B, G) and incidence of metastasis (C, H) and improved overall survival (D, I) and brain metastasis–free survival (E, J) in mice. (K-T) Late Treatment group. (K, P) Representative images showed reduced brain metastasis in CXCR2-IN-1-treated brain metastasis-bearing MDA-IBC3-sEcad (K) and SUM149-sEcad (P). CXCR2-IN-1 reduced the number of brain metastasis lesions (L, Q), incidence of metastasis (M, R) and improved overall survival (N, S) and brain metastasis–free survival (Q, T) in mice. (U, V) Hematoxylin-and-eosin and immunofluorescence stains of brain metastases in the CXCR2-IN-1 Early treatment group (U) and late treatment group (V); CXCR2-IN-1 decreased reactive astrocytes (GFAP+) and CXCR2 in both the MDA-IBC3-sEcad and SUM149-sEcad brain metastases models.

In the ’late treatment’ group, CXCR2-IN-1 significantly reduced brain metastases in both models (Figure 7K, 7P). Analysis of vehicle- and CXCR2-IN-1-treated mice showed a reduction in brain metastasis number (Figure 7L, 7Q) and incidence (Figure 7M, 7R), in addition to prolonged overall survival (Figure 7N, 7S), and brain metastasis–free survival (Figure 7O, 7T). Hematoxylin and eosin staining results for this group are presented in Figure S11E. We evaluated the effect of CXCR2-IN-1 on CXCR2 expression and GFAP+ reactive astrocytes *in vivo* by using hematoxylin and eosin and immunofluorescence staining of brain metastases from vehicle-treated and CXCR2-IN-1-treated groups. In the ’early treatment’ group, CXCR2-IN-1 significantly reduced CXCR2 expression within brain metastases and the surrounding brain parenchyma in both MDA-IBC3-sEcad and SUM149-sEcad models (Figure S11B). The proportions of reactive astrocytes were markedly decreased as well (Figure 7U, Figure S11B). Similar results were observed in the ’late treatment’ group, where CXCR2 expression and GFAP+ reactive astrocytes were reduced in CXCR2-IN-1-treated lesions compared with vehicle-treated controls (Figure 7V, Figure S11C). Together, these findings demonstrate that CXCR2-IN-1 significantly reduces brain metastases and prolongs survival even in established, gross disease, further supporting its potential as an effective therapeutic agent for preventing brain metastases in high-risk patients and treating established brain metastases. These findings suggest that CXCR2-IN-1 modulates the tumor microenvironment by suppressing the sEcad mediated activation of reactive astrocytes.

## DISCUSSION

The molecular mechanisms driving breast cancer cell metastasis to the brain and the interaction of metastases with the brain microenvironment remain poorly understood, which hinders progress in developing new treatments. In this study, we identified sEcad as an indicator of poor-prognosis associated with increased risk of brain metastasis in patients with IBC. By using novel xenograft IBC mouse models, we demonstrated that sEcad is a driver of brain metastasis. sEcad from IBC tumor cells binds to XIAP and activates NFκB signaling to promote cancer cell invasion and anoikis resistance. Tumor secreted sEcad further upregulates CXCR2 in astrocytes which respond to sEcad expressing tumor cells secretion of CXCR2 ligands to promote the metastatic brain niche. Inhibition of CXCR2 blocks sEcad induced pro-metastatic phenotypes and notably, treating sEcad-high brain metastases with a brain-permeable CXCR2 inhibitor reduced metastasis burden and prolonged survival in IBC mouse models. Collectively, our findings highlight the dual role of sEcad in the brain metastatic cascade: (i) promoting invasion and anoikis resistance in breast cancer cells via the XIAP-NFκB axis, and (ii) fostering a supportive brain microenvironment through astrocyte activation and CXCL1/CXCL8-CXCR2 signaling.

One defining feature of metastasis is the dynamic regulation of cadherins, key adhesion molecules that drive cell growth, invasion, and migration. Although E-cadherin is often associated with low metastatic potential,^8,9^ aggressive tumors like IBC paradoxically overexpress it, and its depletion reduces invasion and tumor growth in *in vitro* and *in vivo* models of IBC.^12,18–20^ However, sEcad, the cleaved 80-kDa fragment of E-cadherin, has been linked to tumor progression in several types of cancer and is highly expressed in the serum of cancer patients, correlating with poor prognosis.^36–47^ In breast cancer, high serum sEcad levels are associated with increased metastatic risk and worse outcomes,^68–70^ emphasizing the need to understand its role in metastasis. Prior studies relied on recombinant sEcad protein or neutralizing antibodies, thereby limiting long-term survival and metastasis studies. To address this, we cloned sEcad and generated stable sEcad-expressing cell lines to provide a more robust tool for investigating its function in IBC survival, tumor growth, and brain metastasis. Findings from these sEcad-high stable cell lines mirrored those from recombinant sEcad protein, both demonstrating sEcad’s pro-tumorigenic and pro-metastasis roles in IBC models. Mechanistically, we identified XIAP, a key anti-apoptotic protein, as a novel sEcad binding partner in IBC cells, further supporting sEcad’s role in promoting anoikis resistance. XIAP is a potent inhibitor of apoptosis and enhances the survival of malignant cells.^71–73^ XIAP is also reported to enhance tumor cell invasion and metastasis by driving the pro-survival NFκB pathways.^74–76^ In IBC, XIAP is overexpressed in tumor emboli and is associated with tumor aggressiveness, survival, and chemotherapy resistance.^77–79^ Our findings indicate that sEcad activates XIAP-NFκB signaling, and we reasoned that this activation leads promotion of survival and anoikis resistance, which may explain why aggressive tumors like IBC retain E-cadherin expression.

Notably, while our observations strongly imply that sEcad/E-cadherin directly promote metastasis in IBC, our findings may have broader relevance to other tumor types characterized by elevated levels of these molecules. For example, in prostate cancer, elevated serum sEcad has been associated with advanced metastatic stages, and E-cadherin expression correlates positively with invasive growth and bone metastasis^23–25^. In ovarian cancer, over 80% of tumors display high E-cadherin expression, which is associated with poor patient outcomes and enhanced angiogenesis and invasion of ovarian epithelial cells^26,27^ . In gastric cancer as well, elevated levels of sEcad are associated with increased invasion, and reduced patient survival^28–30^. Similarly, E-cadherin has been linked to poor prognosis in patients with glioblastoma^31^.Collectively, these findings suggest that the oncogenic functions of sEcad/E-cadherin extend beyond IBC to a range of epithelial cancers and taken together with the findings presented here, imply a new mechanism by which E-cadherin promotes aggressiveness across cancers.

While sEcad is traditionally viewed as an extracellular signaling molecule, our findings reveal an intriguing new role. We showed that extracellular sEcad can be internalized by tumor cells and interact with cytoplasmic proteins such as XIAP, a finding which we confirmed using tagged recombinant sEcad.l This is an entirely new finding as our literature search did not reveal any prior demonstration of the ability of extracellular sEcad to enter tumor cells and interact with intracellular, soluble proteins. These results suggest that internalized sEcad has the capacity to modulate intracellular signaling cascades and through them promote tumor cell survival and growth. In this work, we provide evidence of its interaction with XIAP and postulate that this interaction contributes to anoikis resistance in sEcad-overexpressing cells. Supporting the intracellular presence of sEcad, previous studies^27^ in ovarian cancer cells have shown the cytoplasmic expression of sEcad fragments with two potential sources: a membrane-cleaved form and full-length E-cadherin contained within membrane vesicles. Using sucrose gradient centrifugation, they found that while some E-cadherin localized to plasma membrane–enriched fractions, the majority was found in cytoplasmic and Golgi compartments. These findings suggest the existence of multiple sEcad forms - a membrane-cleaved, “internalized sEcad”; “intracellular sEcad” from endogenous processing; and “exosomal sEcad” delivered via vesicles. This complexity reframes sEcad not as a passive byproduct of membrane shedding, but as an active intracellular signaling mediator. Understanding the origin, trafficking, and function of these distinct forms of sEcad will provide important insights into how tumor cells exploit sEcad to promote survival, growth, and metastatic progression.

Examining mechanisms underlying sEcad’s propensity for brain metastasis, we found that sEcad induces reactive astrocytosis in the brain microenvironment by increasing release of pro-inflammatory cytokines from tumor cells, including CXCL1 and CXCL8, and activating their receptor, CXCR2, in astrocytes. In this previously unreported mechanism, sEcad-high cancer cells exploit the CXCL1/CXCL8-CXCR2 axis to create a supportive niche for brain metastasis growth. CXCR2 is involved in various neurological conditions, including neurological repair in multiple sclerosis, optic nerve injury, and neuropathic pain; it is also expressed in immune cells, endothelial cells, and mesenchymal stem cells.^80–86^ In brain injury and autoimmune disease models, CXCR2 participates in maintaining BBB integrity,^87,88^ whereas in brain metastasis it has been shown to reprogram neutrophils to promote tumor progression.^89^ Given the growing evidence that reactive astrocytes foster brain metastasis by compromising the BBB^59–64^ thus allowing easier access for tumor cells, we investigated whether sEcad-mediated brain metastasis and astrocyte activation is driven by CXCR2. To the best of our knowledge, no prior studies have explored CXCR2-mediated astrocyte activation as a mechanism for brain metastasis, and our findings strongly support this hypothesis through both *in vitro* studies and IBC brain metastasis models. We further demonstrate that the CXCR2 antagonist CXCR2-IN-1 effectively blocks brain metastasis and prolongs survival in preclinical mouse models, highlighting inhibition of CXCR2 as a potential therapeutic strategy that should be more extensively investigated in preclinical and clinical studies on metastatic brain progression.

In summary, this study reveals sEcad as a key driver of brain metastasis in IBC, demonstrating its critical role in several steps in the metastatic cascade. We demonstrate that sEcad not only enhances tumor cell-intrinsic survival but also reshapes the brain microenvironment by activating astrocytes, fostering a pro-metastatic niche that drives selective brain colonization. Our findings that we can achieve inhibition of brain metastasis with a BBB-penetrant inhibitor of CXCR2 highlight the therapeutic potential of targeting the sEcad-CXCL1-CXCR2 pathway in aggressive cancers. Disrupting this axis is a promising strategy to limit brain metastasis in IBC and other malignancies that metastasize to the brain.

## RESOURCE AVAILABILITY

### [Lead contact]

Further information and requests for resources and reagents should be directed to and will be fulfilled by the lead contact, Bisrat G Debeb.

### [Materials availability]

Plasmids generated in this study are available upon reasonable request from the corresponding author.

### [Data and code availability]

The mass spectrometry proteomics data have been deposited to the ProteomeXchange Consortium via the PRIDE partner repository with the dataset identifier PXD063581.

## ACKNOWLEDGEMENTS

We thank Alma Faust, PhD, for critical review of the manuscript and helpful discussions and Christine F. Wogan, MS, ELS, of MD Anderson’s Division of Radiation Oncology for scientific review and editing of the manuscript. We thank Jordan Pietz, MA, CMI from the MD Anderson Biomedical Visualization team for his assistance in creating the visual artwork. We thank Research Histology Core Laboratory (supported by NIH/NCI Core Grant P30CA016672) at MD Anderson for immunohistochemical analysis. We thank the Flow Cytometry and Cellular Imaging Core Facility(supported by NIH/NCI Core Grant P30 CA016672) at MD Anderson Cancer Center for confocal microscopy and technical support with flow sorting. We thank Dr. Funda-Meric Bernstam for generously providing us the BCX010 cell line. We thank Dr. Clark D Wells (Indiana University) for generously providing the normal human astrocyte (NHA) cells. This work was supported by grants from DoD BC230452 (BGD), METAvivor (BGD), NIH/NCI 1R01CA284102 (BGD/WAW/MR), ACS RSG-19126–01 (BGD), MD Anderson Cancer Center (Bridge Funding, Startup, and Bootwalk funds), and the State of Texas Grant for Rare and Aggressive Cancers.

## Contributors: MDACC Inflammatory Breast Cancer Team Authorship

The prospective IBC annotated biobank exists through the collaborative efforts of the following individuals who manage the protocol and regulatory compliance, regularly meet to review and adjudicate inconclusive diagnoses, accrue and consent patients, coordinate sample collections, maintain records, review data collection and specimen inventory, and provide researchers access to the materials through secondary IRB approved protocols.(Saleem,Sadia; Valero,Vicente; Lim,Bora; Nasrazadani,Azadeh; Layman,Rachel M;Lucci,Anthony; Sun,Susie Xinying; Woodward,Wendy;Stauder,Michael C; Whitman,Gary J; Le-Petross,Huong; Patel,Miral Mahesh; Lu,Yang; Marx,Angela N; Alexander,Angela; Yajima,Chasity L; Kai,Megumi; Villarreal,Lily A; Lopez,Heather B;)

## Author contributions

X.H. and B.G.D. conceived and designed the project. X.H. performed most of the experiments, analyzed the data, and interpreted the results., Y.X and H.Z were performed the TAP-MS and BioID experiments and analysis. J.S. provided statistical analysis support. S.K. provided pathological expertise and analysis of xenograft tumors. E.S.V, J.C, M.R, N.F, C.B, D.T, and W.A.W. provided resources and contributed to revision of the manuscript. X.H. and B.G.D. wrote and edited the manuscript with input from all other authors. B.G.D. supervised the study.

## STAR METHODS

Key resources table.

## EXPERIMENTAL MODEL AND STUDY PARTICIPANT DETAILS

### Plasmid construction

The soluble E-cadherin cDNA (1-707aa) was amplified from E-cadherin (Gene ID: 999) cDNA and then cloned into modified LentiV_Blast-Flag (#111887, Addgene), pHA-N1 (#86049, Addgene), pcDNA3-neo-Strep-flag-Cterm (#102645, Addgene), and MCS-BioID2-HA (#74224, Addgene) vectors respectively. The pEBB-HA-XIAP (#11558), pEBB-HA-XIAP-BIR2(#25688), pEBB-HA-XIAP-BIR3(#25689) and pEBB-HA-XIAP-BIR1-2-3(#25692) plasmids were purchased from Addgene.

### Lentiviral production and transduction

Protocol details are described elsewhere.^12,51^ Briefly, Lipofectamine 3000 (Invitrogen, USA) DNA mixture (10 µg LentiV_Blast-Flag-sEcadherin/Luc-GFP, 7.5 µg of psPAX2 packaging plasmid and 2.5 µg of pMD2.G enveloping plasmid) were incubated overnight into HEK293T cells. The culture medium was then removed and replaced with fresh medium. The supernatant containing the virus was collected, filtered through a 0.45-μm HV Durapore membrane (EMD Millipore) to remove cells and large debris, and concentrated by ultracentrifugation. Target cells with a confluence of about 70% were used for transduction. The medium was changed 2 hours before transduction. Lentiviral transduction was performed in the presence of 8 μg/mL polybrene (Sigma-Aldrich, USA) for 24 hours, followed by replacement with fresh medium.

### Antibodies and reagents

Anti-GAPDH (#8884), Anti-β-Actin (#4967), Anti-Tubulin (#2148S), Anti-HA-Tag (#C29F4) (3724S), Anti-XIAP (#14334S), Anti-NF-κB p65 (#8242), Anti-Phospho-NF-κB p65 (Ser536) (#3033), Anti-Caspase-3 (#9662), Anti-Caspase7 (#9494), Anti-Cleaved Caspase-9 (Asp353) (#9509) , Anti-GFAP (#3670S), and Anti-Ki67(#12202) antibodies were purchased from Cell Signaling Technology (Beverly, MA, USA). All the secondary antibodies [Anti-mouse-HRP (#7076S), Anti-rabbit-HRP (#7074S), Anti-mouse-Alexa Fluor® 555 (#4409S), Anti-mouse-Alexa Fluor® 488 (#4408S), Anti-rabbit-Alexa Fluor® 594 (#8889S), and Anti-rabbit-Alexa Fluor® 488 (#4412S)] were also purchased from Cell Signaling Technology. Anti-CXCR2 (ab65968) was purchased from Abcam (Cambridge, MA, USA). High-capacity streptavidin agarose (#20359), S-protein agarose (#69704) and Protein A/G agarose (#20422) were purchased from Thermo Scientific (Rockford, IL, USA). Human E-Cadherin Antibody (MAB1838), Recombinant Human E-Cadherin Protein-CF (sEcad protein) (8505-EC) , Human E-Cadherin Quantikine ELISA Kit (#DCADE0B), Human CXCL1/GRO alpha Quantikine ELISA Kit (#DGR00B), Human IL-8/CXCL8 Quantikine ELISA Kit (#D8000C), Human Dkk-1 Quantikine ELISA Kit (#DKK100B), Proteome Profiler Human Cytokine Array Kit (#ARY005B), and Proteome Profiler Human Apoptosis Array Kit (ARY009) were purchased from R&D system (USA). Anti-Flag (#F1804), RIPA buffer (#R0278), and sEcad neutralizing antibody DECMA1 (#MABT26) were purchased from Sigma (St Louis, MO, USA). SB225002 (#S7651) was purchased from Selleck (Houston, TX, USA). CXCR2-IN-1 was purchased from MedChem Express (#HY-101022).

### Cell cultures

SUM149,and SUM190 cell line was purchased from Asterand (Detroit, MI), and the MDA-IBC3 cell line was generated in the laboratory of Dr. Wendy Woodward at MD Anderson.^53,90^ The human TNBC cell line BCX010, derived from a patient with triple-negative inflammatory breast cancer, was generously donated by Dr. Funda Meric-Bernstam (MD Anderson Cancer Center)^91^ .The cell culture conditions were described previously.^12,92^ HEK293T cells were obtained from the American Type Culture Collection (Manassas, VA, USA) and were cultured in Dulbecco’s modified Eagle’s medium (DMEM) supplemented with 10% FBS and 1% penicillin and streptomycin (#15140122, Invitrogen, Carlsbad, CA, USA) at 37oC in a humidified incubator with 5% CO2. Normal human astrocytes were cultured with High DMEM (HyClone #SH30243.01 GE),10% fetal bovine serum (FBS) (Gibco #16000-044), 1% Pen/Strepto (#15-140-122, Fisher Scientific), 5μg/mL Insulin (Cell Applications #128-100), 10 μM Hydrocortisone (#H0888, Sigma), and 5 μg/mL N-acetylcysteine (A9165, Sigma). The human microvascular endothelial cell line HBEC-5i (CRL-3245), mouse astrocyte C8-D1A (CRL-2541), human umbilical vein endothelial cells (HUVECs) (PCS-100-010), and mouse endothelial cells bEnd.3 (CRL-2299) cells were acquired from the American Type Culture Collection (ATCC) (Manassas, VA, USA) and cultured per the recommended protocols. Bend3 endothelial cells and C8-D1A astrocytes were cultured in ATCC-formulated Dulbecco’s Modified Eagle’s Medium (30-2002, ATCC) containing 10% FBS (30-2020, ATCC). HMC3 cells were cultured in EMEM (30-2003, ATCC) basal medium containing 10% FBS (30-2020, ATCC). HUVECs were cultured with Vascular Cell Basal Medium (PCS-100-030, ATCC), Endothelial Cell Growth Kit-VEGF (PCS-100-041, ATCC), and Penicillin-Streptomycin-Amphotericin B Solution (PCS-999-002, ATCC).

### Enzyme-linked immunosorbent assay of IBC samples

Serum samples were identified from a prospectively maintained IBC registry and biobank. All patients provided informed consent for collection and use of the samples on an IRB approved protocol. Levels of soluble E-cadherin in serum from 348 patients with IBC were analyzed by ELISA (R&D Systems, Minneapolis, MN, USA #DCADE0) according to the manufacturer’s instructions. Samples were assayed in duplicate. For conditioned-medium ELISA, cancer cells were cultured with serum-free medium for 36 h, after which the conditioned medium was collected and subjected to ELISA for CXCL1, CXCL8 and DKK1 (R&D Systems, Minneapolis, MN, USA #DGR00B, #D8000C, and #DKK100B, respectively).

### Western blotting

Total protein was extracted from cancer cells using RIPA buffer (Sigma) with 10 µL/mL phosphatase and 10 µL/mL protease inhibitor cocktail. Forty micrograms of protein lysate from each sample were electrophoretically separated with a 10% SDS–PAGE gel and then transferred to a polyvinylidene difluoride membrane. SDS-PAGE and immunoblotting were done as previously described by us.^12,51^ Membranes were incubated with the corresponding primary antibodies overnight at 4°C and then incubated with secondary antibodies (1:5000) anti-rat IgG (#HAF005, R&D Systems) and anti-rabbit IgG (#7074, Cell Signaling) for 2 h at room temperature. GAPDH/B-actin/Vinculin were used as an internal control.

### Xenograft studies

Animal experiments were done in accordance with protocols approved by the Institutional Animal Care and Use Committee of MD Anderson Cancer Center, and mice were euthanized when they met the institutional euthanasia criteria for tumor size and overall health condition. Animal care and use were in accordance with institutional and NIH guidelines. For *in vivo* brain metastasis studies, control and sEcad-overexpressing SUM149 cells (2x10^5^) were injected intracardially and MDA-IBC3 cells (1x10^6^) were injected into the lateral tail veins, into 4-to 6-week-old female SCID/Beige mice (n = 10 per group) (purchased from Harlan Laboratories, Indianapolis, IN, USA). In brain metastasis studies, mice received daily CXCR2-IN-1 (2 mg/kg) treatment via oral gavage. For the ’Early treatment’ study, treatment was initiated a day before the injection of sEcad cancer cells. For the ’Late treatment’ studies, treatment was initiated 4 weeks after cancer cell injection.

### Endothelial cell adhesion assay

Human brain microvascular endothelial HBEC-5i cells were seeded into 6-well plates and allowed to form a confluent monolayer for 24 hours. The tops of the monolayers were then seeded with sEcad overexpressing or control MDA-IBC3 / SUM149 cells labeled with GFP. The cells were allowed to incubate for 30 minutes, media was aspirated, and cells were washed with PBS twice to remove nonadherent cells. The fluorescent tumor cells were imaged, and the numbers of cells were counted per field as described previously.^93^

### *In vitro* trans–blood brain barrier migration assay

Mouse astrocytes C8-D1A (5× 10^5^) were plated on the bottom side of a transwell, and cell medium was refreshed every 15 mins for 6 hours. The transwell was then inverted back and 2.5× 10^5^ bEnd.3 cells were plated on the top side of the membrane, after which the transwell was incubated at 37°C for 3 days to allow formation of blood-brain barrier (BBB). Transwells with intact BBB were washed twice with phosphate-buffered saline (PBS) to remove the serum and inserted into 24-well plates containing the tumor-conditioned medium from the tumor cells in the bottom chamber after the cells had been starved for 24 hours. Green fluorescence protein (GFP)-labeled tumor cells were then plated onto the top chamber and the migratory ability of the tumor cells was noted 20 hours later.^93^

### *In Vitro* angiogenesis assay

About 2× 10^4^ HUVECs were plated on Extracellular Matrix Gel provided in an *in vitro* angiogenesis kit (Abcam, ab204726) with tumor-conditioned medium and monitored for tube formation up to 18 hours. For treatment, 20 μg/mL sEcad recombinant protein, 20 μg/mL DECMA1 neutralizing antibody, and IgG control were added to the fresh HUVEC medium for 24hours. Branch points were quantified per microscopic field.

### Proteome profiler arrays

Human apoptosis array (#ARY009) and cytokine array (#ARY005B) kits (R&D Systems) were used according to the manufacturer’s instructions. Briefly, MDA-IBC3-Con/sEcad and SUM149-Con/sEcad cells were serum-starved for 24 hours and fresh growth medium was added. To prepare samples for apoptosis array, 400 μg of cell lysate pooled from three independent replicates were used for each sample. To prepare samples for cytokine array, 700 μL of cell culture medium pooled from three independent replicates were used. The stained arrays were imaged with the Bio-Rad gel documentation system and quantified with ImageJ.

### Conditioned medium

To collect conditioned medium, cancer cells were seeded into culture dishes and incubated for 24 hours. After, cells were washed twice with 1X PBS, 10 mL of fresh medium without FBS was added and the cells incubated for another 36 hours at 37°C, and then the medium was collected and filtered with a 0.2-μm syringe filter. For cytokine assays, the amounts used were described in the manufacturer’s protocol. For astrocyte co-culture experiments, the cells were washed twice with 1x PBS after 24 hours of attachment, and then treated with 30% conditioned medium and 70% culture medium for 24 hours to collect samples for western blot assays.

### Mass spectrometry sample preparation and analysis

Proteins for affinity purification for mass spectrometry (MS) were obtained as described previously.^51^ Briefly, pellets of HEK293T cells expressing Sbp-flag-sEcad and BioID2-tagged-sEcad were harvested and washed with 1X PBS. Cell pellets were lysed with 1X NETN buffer (100 mM NaCl, 1 mM EDTA, 20 mM Tris-HCl, and 0.5% Nonidet P-40) with a protease inhibitor cocktail (Sigma-Aldrich) for 20 min at 4°C. After centrifugation at 16,000 × *g* for 20 min at 4°C, the supernatant was incubated with streptavidin-conjugated beads (Thermos Fisher Scientific) at 4°C for 4 hours. The beads were then washed four times with NETN buffer. Samples were subjected to SDS–PAGE, after which the gel was fixed and stained with Coomassie brilliant blue (# 1610436, Bio-Rad). The entire lane of the sample in the gel was then excised and subjected to MS analysis.

For the MS, gel pieces were de-stained twice with 50% acetonitrile in 50 mM ammonium bicarbonate, followed by dehydration with acetonitrile. The pieces were digested overnight at 37°C with trypsin (V5280, Promega) in 25 mM ammonium bicarbonate. Digested peptides were extracted with acetonitrile and dried by vacuum. Dried samples were reconstituted in 0.1% formic acid and analyzed by using a 75-minute gradient at a flow rate of 300 nL/min (solvent A: 0.1% formic acid in water; solvent B: 0.1% formic acid in acetonitrile). The gradient program was as follows: 5% to 24% solvent B over 60 minutes, 24% to 45% solvent B from 60 to 66 minutes, 45% to 90% solvent B from 66 to 69 minutes, and a hold at 90% solvent B from 69 to 75 minutes. The eluate was analyzed by using a Q Exactive HF-X Hybrid Quadrupole-Orbitrap mass spectrometer (Thermo Fisher Scientific) in data-dependent mode. Precursor MS spectra were acquired in the 375-1300 m/z range with a resolution of 120,000. The top 40 ions were selected for fragmentation via collision-induced dissociation at a normalized collision energy of 28, with an isolation width of 1.2 Da, a dynamic exclusion time of 20 seconds, an AGC target of 1× 10^5^, and a maximum injection time of 60 ms.

For MS data analysis, raw data were processed by using MaxQuant software (version 1.6.6.0; Max Planck Institute of Biochemistry) with a false discovery rate (FDR) <0.01 at the protein and peptide-spectrum match levels. The Human UniProt FASTA database (July 2019, containing 27,183 entries) was used. Acetylation of protein N-termini and oxidation of methionine were set as variable modifications. Trypsin was specified as the enzyme, and a maximum of two missed cleavages was allowed. Precursor mass tolerance was set to 10 ppm, with fragment mass tolerance at 0.01 Da. MS results were compared with background HEK293T proteome profiling data.^94^ Proteins with a fold-enrichment above the average were identified as candidate binding proteins.

### Immunoprecipitation

Immunoprecipitation was done as described previously.^12,51^ Briefly, HEK293T cells were seeded 24 hours before transfection. Cells were transiently transfected with Lipofectamine 3000 (#L3000015, Invitrogen, USA), collected 24 hours later, and then lysed in 1X RIPA buffer (#R0278, Sigma-Aldrich, USA) containing 10 µL/mL phosphatase inhibitor cocktail and 10 µL/mL protease inhibitor cocktail (Santa Cruz Biotechnology, USA) for 10 minutes. For exogenous immunoprecipitation with streptavidin beads, suspensions of streptavidin beads and cell lysates were incubated for 2.5 hours on an orbital shaker at 4°C, washed 5 times with 1X RIPA buffer, and the pellets were resuspended in SDS sample buffer and boiled for 8 minutes. For endogenous immunoprecipitation, HEK293T cell lysates were incubated with 25 μL protein A/G beads with anti-Flag or anti-XIAP antibody, and the mixtures were incubated for 4 hours on an orbital shaker at 4°C. After centrifugation, the pellets were washed five times in 1X RIPA buffer, resuspended in 1X SDS sample buffer, and boiled for 8 minutes.^12,51^

### Immunohistochemical and immunofluorescence staining

Details of the immunofluorescence staining protocol for cultured cells was described by us previously.^12^ Astrocytes were grown on Millicell EZ SLIDE 4-well glass (PEZGS0416) and treated with the corresponding proteins or inhibitors. Cells were then stained with the primary antibodies anti-GFAP (1:200 dilution, CST, #3670S) and anti-CXCR2 (1:100 dilution, Abcam, #ab65968) for 2 hours at room temperature. Staining was visualized by using Alexa-Fluor 594-conjugated secondary antibody (diluted 1:500) and Alexa-Fluor 468-conjugated secondary antibody (diluted 1:500) and then stained with 4’6-diamidino-2-phenylindole (DAPI) with indole (0.5 μg/mL) for 15 minutes to label the nuclei. Finally, the coverslips were washed five times with 1X PBS. Immunofluorescence microscopy images were obtained with a Keyence BZ-X800 microscope (Keyence Corporation of America).

Formalin-fixed, paraffin-embedded sections of brain tissues were stained with hematoxylin and eosin (H&E) at MD Anderson’s Pathology core facility with standard and validated protocols. Slides were analyzed by a pathologist specializing in breast cancer (SK). Immunohistochemical staining was done with biotinylated secondary antibodies (Vector), biotin-avidin-peroxidase complex (Vector), and diaminobenzidine (brown; Sigma) or Vector blue (Vector) as the developing agents. Fluorescence immunohistochemical staining was done with the Alexa Fluor-tagged secondary antibodies Alexa 488 (green) or Alexa 594 (red). Primary antibodies used were rabbit anti–CXCR2 (1:100), mouse anti-GFAP (1:500), and DAPI (Molecular Probes, D-1306) was used as a fluorescent counterstain. Stained sections were examined and photographed with bright-field and fluorescence microscopy with a Keyence BZ-X800 microscope (Keyence Corporation of America).

### Soft agar / anchorage-independent growth assay

Cell growth in soft agar for anchorage-independent growth were assessed as described elsewhere.^51,95^ Briefly, 1 mL of complete medium containing 1% agarose was added evenly to each well of a 12-well plate, while ensuring no bubbles were present, until it completely solidified (bottom layer). MDA-IBC3 cells or SUM190 (5000 cells each) or SUM149 cells or BC0X10 cells (8000 cells each) were suspended in 0.5% agarose in complete medium in the presence or absence of sEcad recombinant protein (20 μg/mL) and DECMA1 (20 μg/mL) (top layer). Then, the top layer mixture of cells and agarose was layered onto the solidified bottom layer and the layers were incubates at 37°C for 3 weeks. Colonies were evaluated with an inverted microscope (Nikon, Tokyo, Japan) and 10 random areas were chosen to observe and photograph. In addition, colonies were stained with MTT, and colonies >80 µm in diameter were counted with the GelCount system (Oxford Optronix Ltd). The same experiments and analyses were done with control and sEcad-overexpressing MDA-IBC3, SUM190, SUM149, or BCX010 cells.

### Anoikis assay

The anoikis assay was performed as described previously.^51^ Cells at high density (1 × 10^6^/mL) were harvested by trypsinization and suspended in tissue culture plates coated with poly-HEMA (10 mg/mL; Sigma). Cells were harvested 24 hours later, washed with 1X PBS, stained with an FITC-conjugated Annexin V apoptosis detection kit (BD Biosciences, Franklin Lakes, NJ, USA) according to the manufacturer’s protocol, and analyzed by flow cytometry.

### *In vitro* migration and invasion assays

Migration assays were done with 24-well transwell plates (Corning, Inc. USA). For invasion assays, the upper chamber was pre-coated with Matrigel (BD Biosciences, Franklin Lakes, NJ, USA). To investigate the effects of sEcad overexpression on IBC cell migration and invasion, 5 × 10^4^ Control or sEcad-overexpressing SUM149 or BCX010 cells were placed in 100 μL of serum-free medium and seeded into the upper chamber with or without Matrigel. IgG, 20 μg/mL sEcad-recombinant protein, or 20 μg/mL DECMA1 neutralizing antibody were added to the upper chamber containing 5 × 10^4^ SUM149 or BCX010 cells, and migration and invasion were analyzed as follows. After 24 hours of culture, cells that had migrated to the bottom surface were fixed in 4% paraformaldehyde (#AAJ19943K2, Thermo Scientific, USA) for 30 minutes and stained with 1% crystal violet solution for 25 minutes, and the non-migrated cells were gently removed from the upper chamber with a cotton swab. Under a microscope (Nikon eclipse Ti camera, NY, USA), ten randomly chosen visual fields were recorded and analyzed with Image J software.

### Statistical analysis

Patient characteristics were summarized by serum sEcad levels from patients with IBC compared between patients with sEcad-Low (<95 ng/mL) or sEcad-High (>95 ng/mL) levels (99.8 ng/mL was the 90th percentile). Univariate and multivariate Cox regression analyses were used for overall survival and breast cancer–specific survival as outcome variables. Univariate and multivariate Fine-Gray models were used to consider time to metastasis and time to brain metastasis as outcome variables, and death without an event of interest was considered a competing risk event in the Fine-Gray model. The Kaplan-Meier method was used to estimate survival probabilities, and cumulative incidence was estimated by the Aalen-Johansen method. Log-rank tests were used to compare survival curves. The Gray test was used to compare cumulative incidence curves. *P* values of <0.05 indicated statistically significant differences. SAS 9.4 (SAS institute INC, Cary, NC) was used for data analysis. All *in vitro* experiments were repeated at least three times, and graphs depict mean ± SEM. Statistical significance was determined with Student’s *t* tests (unpaired, two-tailed) unless otherwise specified. GraphPad software (GraphPad Prism 8, La Jolla, CA) was used.

## SUPPLEMENTARY INFORMATION

Supplementary Figures S1-S11

Table1-2

