## Supplementary Figures S1-S11 for "Soluble E-cadherin Drives Brain Metastasis in Inflammatory Breast Cancer"

**SUPPLEMENTARY INFORMATION**

**Supplementary Figures S1-S11**

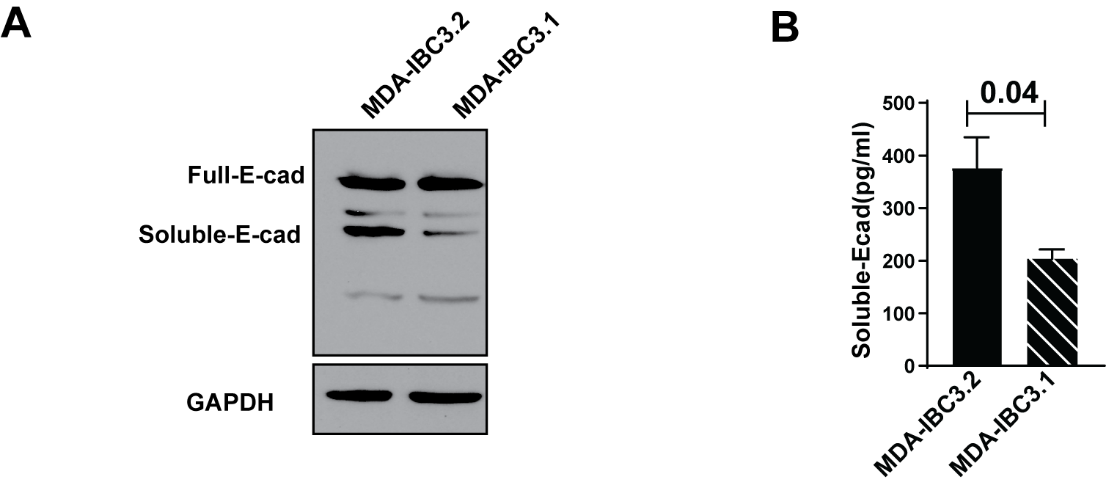

**Figure S1. sEcad expression levels are higher in brain-metastasizing cells.**

(A, B) Western blot and enzyme-linked immunosorbent assay findings show higher expression of sEcad in the highly brain metastasizing MDA-IBC3.2 cells compared to weakly brain metastatic MDA-IBC3.1 cells.

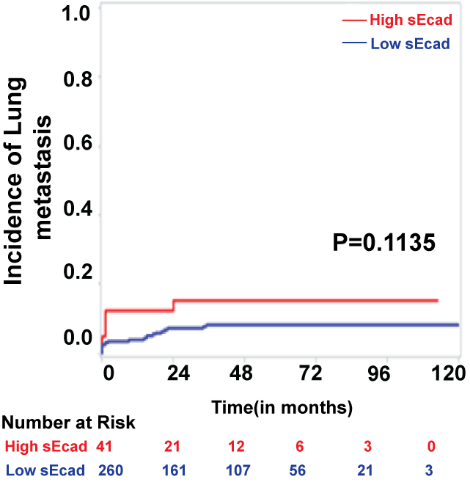

**Figure S2. Serum sEcad levels are not correlated with the development of lung metastasis in patients with inflammatory breast cancer.**

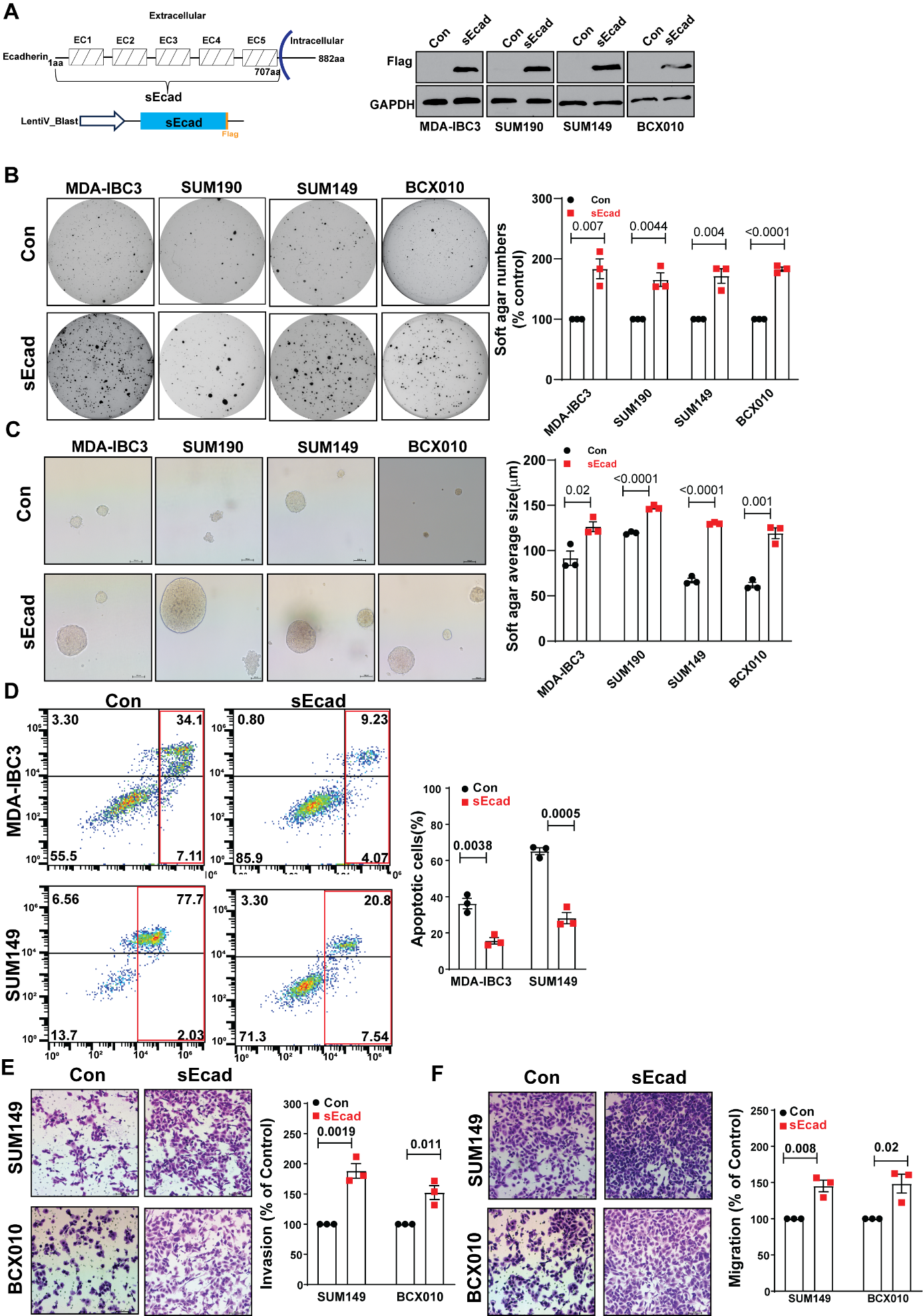

**Figure S3 [previous page]. Ectopic overexpression of sEcad promotes anchorage-independent growth, migration, invasion, and anoikis resistance in IBC cells in vitro.**

(A) Generation of sEcad-FLAG overexpressing IBC stable cell lines. The total cell lysates of the 4 IBC cell lines (MDA-IBC3, SUM190, SUM149, BCX010) were analyzed by western blot with anti-FLAG and anti-GAPDH (internal control) antibody.

(B, C) sEcad overexpression increases the number (B) and size (C) of soft gar colonies in IBC cells

(D) sEcad overexpression inhibits anoikis in IBC cells. Control or sEcad over-expression MDA-IBC3 and SUM149 cells were treated with poly-HEMA. 24h later, cells were harvested and analyzed by flow cytometry using FITC-Annexin V/PI- kit. The representative FACS analysis were shown (Right) with quantitation of three independent experiments (Left).

(E) sEcad promotes migration of SUM149 and BCX010 IBC cells.

(F) sEcad promotes invasion of SUM149 and BCX010 IBC cells.

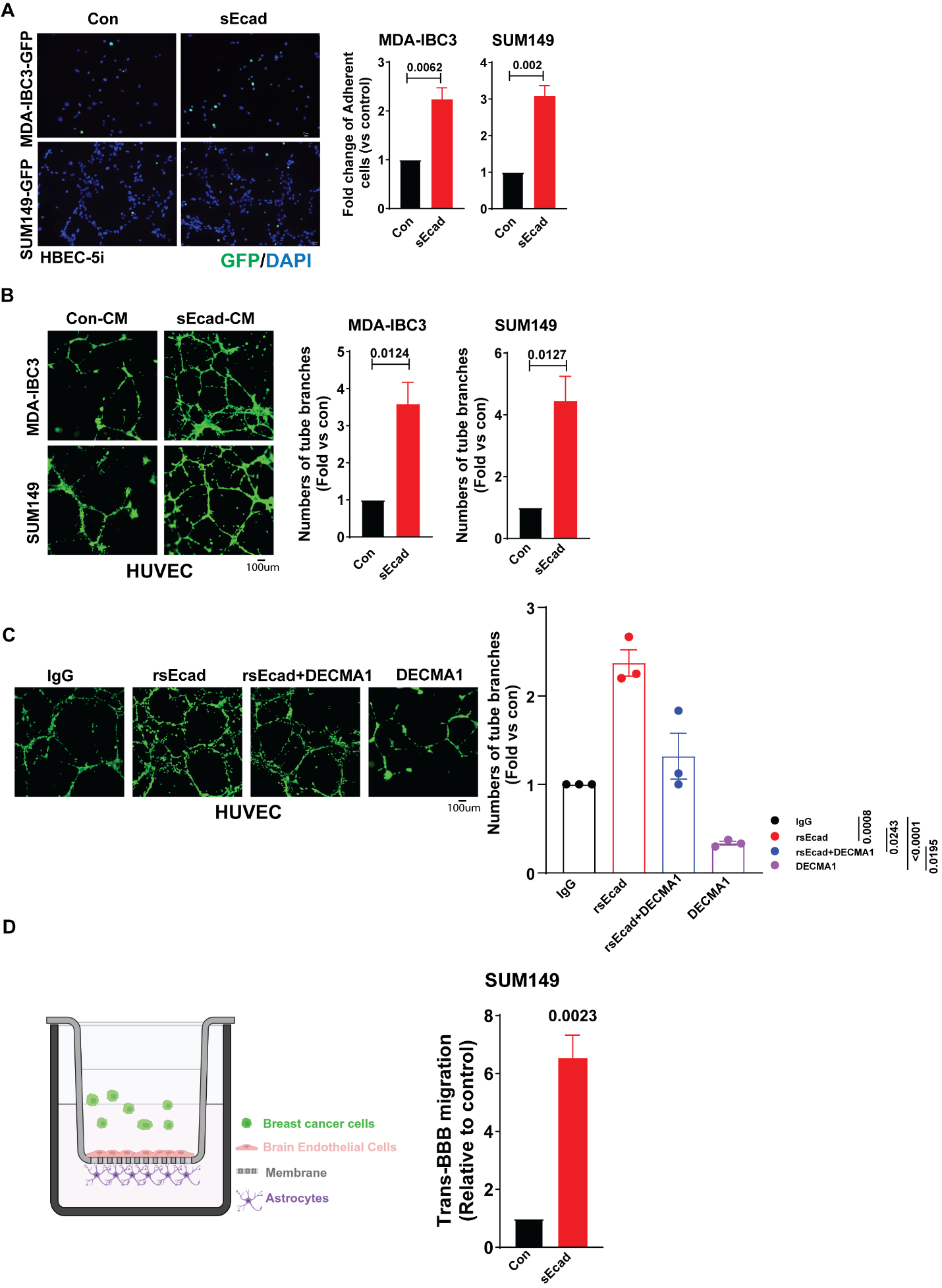

**Figure S4 [previous page]. sEcad promotes in vitro endothelial cell adhesion, angiogenesis and trans-endothelial migration.**

(A) sEcad promotes brain endothelial cell adhesion. GFP-labeled sEcad-overexpressing IBC cells showed enhanced adhesion to human brain microvascular endothelial cells (HBEC-5i) *in vitro* compared with control cells. Data are represented as mean ± SEM from at least three independent experiments.

(B) *In vitro* angiogenesis assay of human umbilical vein endothelial cells (HUVECs) treated with conditioned medium from MDA-IBC3 or SUM149 control and sEcad-overexpressing cells. Data on the right represent mean ± SEM of three biological replicates.

(C) sEcad protein promotes angiogenesis, an effect countered by DECMA1. Quantification data represent mean ± SEM of three biological replicates.

(D) In vitro BBB trans-endothelial migration model showed sEcad-overexpressing SUM149 cells had a higher migrating ability across the trans-BBB assay (left). Quantitation of trans-BBB migrating cells is shown on the right.

**Figure S5: sEcad gets internalized into IBC tumor cells.**

(A) IF staining visualizes cytoplasmic localization of sEcad (green) in SUM149 and MDA-IBC3 cells; HA-sEcad was transfected into MDA-IBC3 and SUM149 cells and the subcellular localization of HA-sEcad by using anti-HA polyclonal antibody. Representative images are shown (60x magnification);

(B)IF shows the internalization of sEcad (green; FITC-tagged anti-His) in MDA-IBC3 and SUM149 cells treated with recombinant sEcad protein (20ug/ml, 36hrs). Representative images are shown (20x magnification).

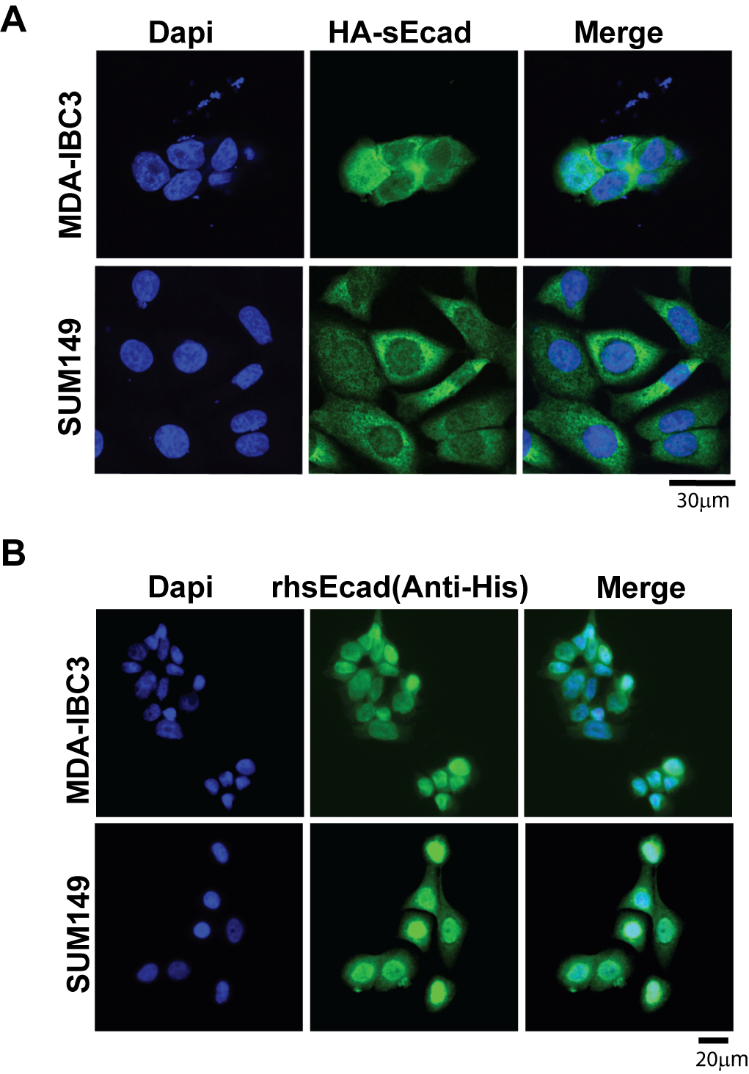

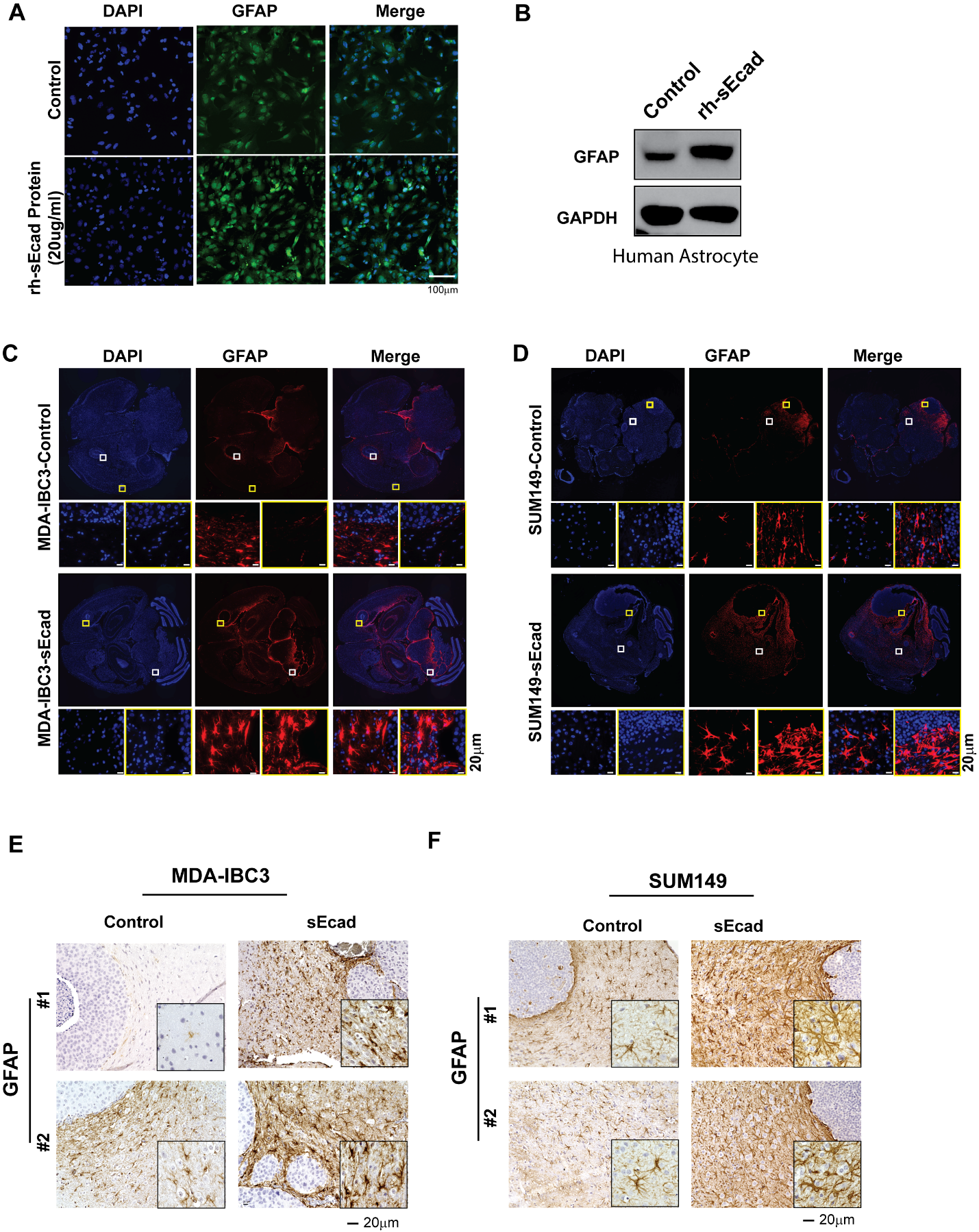

**Figure S6 [previous page]. sEcad induces reactive astrocytes**

**(**A) and (B) Astrocytes treated in vitro with sEcad recombinant protein (20 µg/ml for 36 h) show increased GFAP+ reactive astrocytes by immunofluorescence (IF) (A) and immunoblotting (B); (C) and (D) IF staining shows that metastatic lesions from sEcad-expressing tumors have significantly higher GFAP+ reactive astrocytes vs the control group. Data shown are from MDA-IBC3 tail-vein injection (C) or intracardiac injection (D) experiments.

(E) and (F) Immunohistochemical stains of brain metastases generated from MDA-IBC3 (E) and SUM149 (F) control and sEcad-overexpressing cells confirm the upregulated expression of GFAP in the sEcad-overexpressing cells in mouse xenograft brain metastatic lesions.

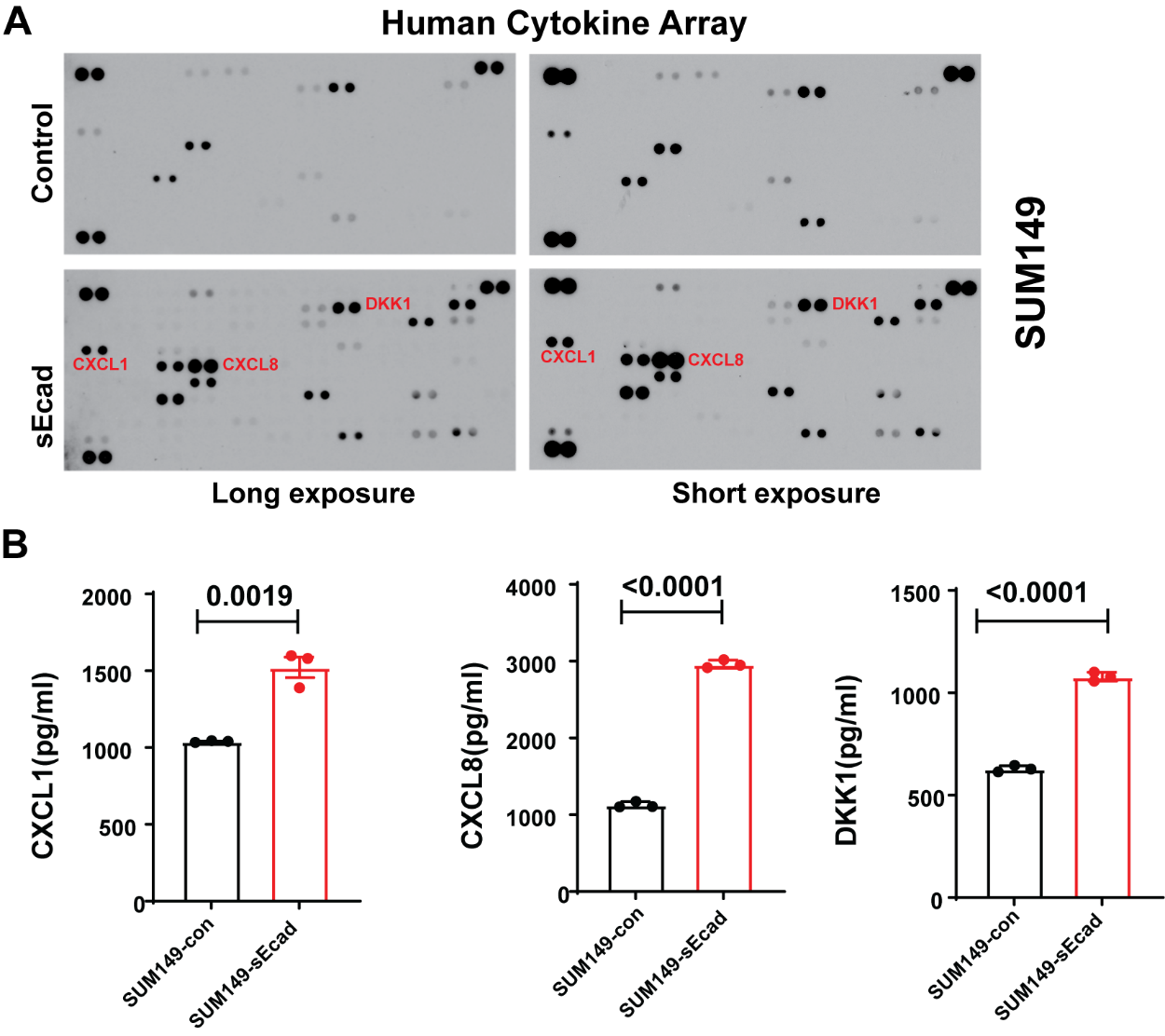

**Figure S7. Conditioned medium from SUM149 sEcad-overexpressing cells induces CXCL1/CXCL8/DKK1 levels.**

(A, B) Human cytokine array analysis of secreted factors from sEcad-overexpressing SUM149 cells. (A) Increased levels of cytokines including CXCL1, CXCL8 and DKK1 in conditioned medium from sEcad-overexpressing SUM149 cells. (B) Validation of the levels of detected cytokines from (A) by enzyme-linked immunosorbent assay.

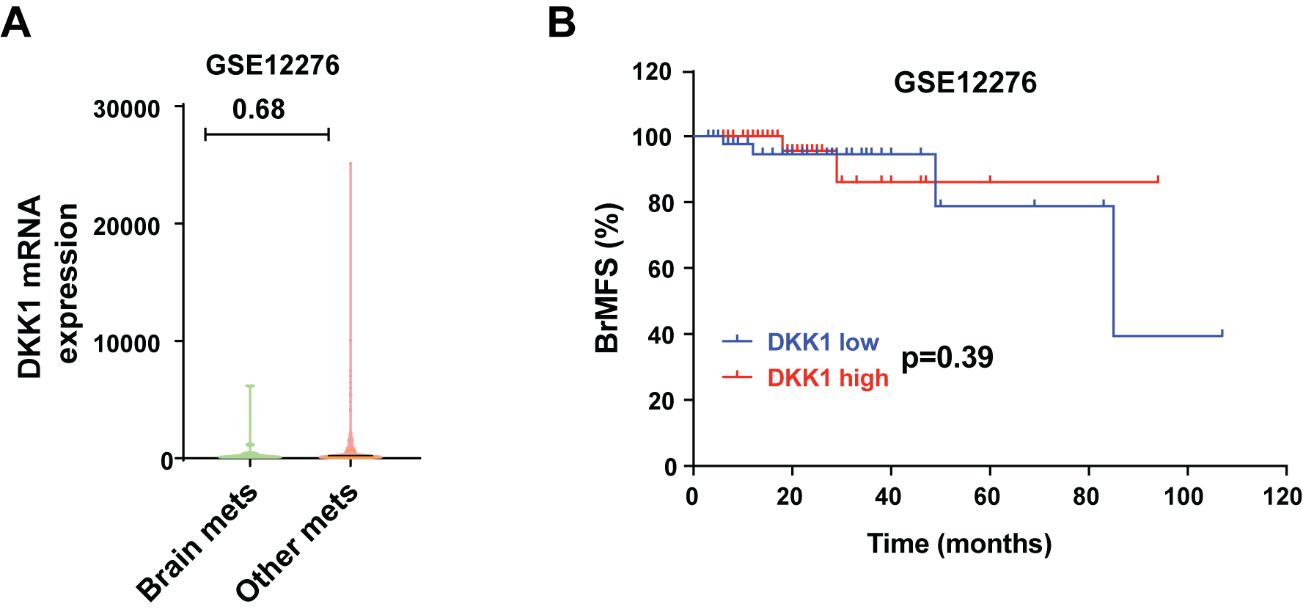

**Figure S8. DKK1 was not correlated with brain metastasis–free survival.**

(A) DKK1 expression did not differ between patients with brain metastases and patients with metastases at other sites in the Gene Expression Omnibus (GEO) database GSE12276.

(B) DKK1 expression levels were not significantly correlated with brain metastasis-free survival (BrMFS), (GSE12276); low and high indicate the bottom tertile (25th) and top tertile (75th), respectively.

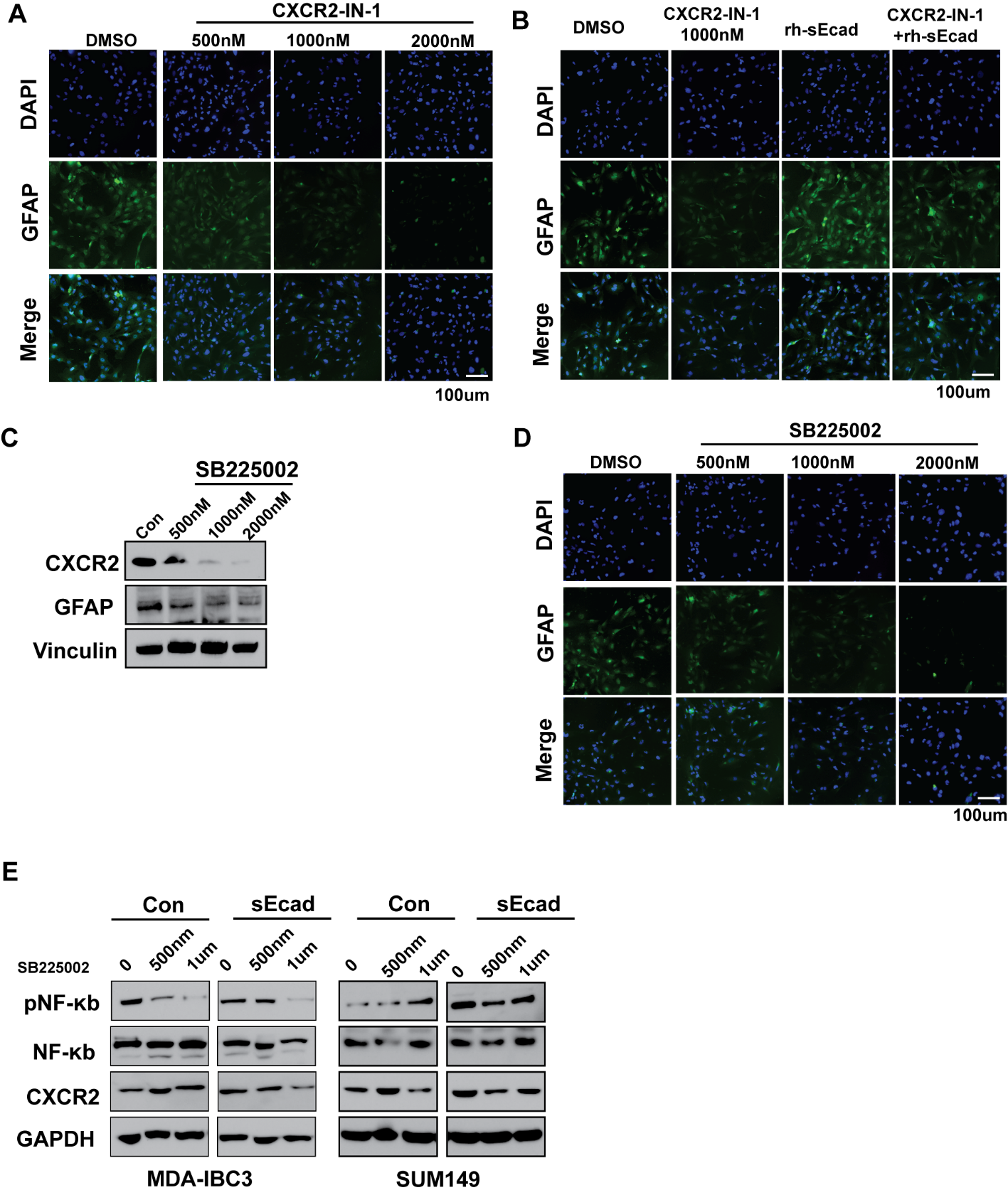

**Figure S9 [previous page]. Blockade of CXCR2 reduces reactive astrocytes in vitro.**

(A) Immunofluorescence staining shows that CXCR2-IN-1 reduced the numbers of GFAP+ astrocytes in vitro.

(B The CXCR2 inhibitor CXCR2-IN-1 can inhibit GFAP+ astrocytes induced by recombinant sEcad protein in vitro.

(C, D) The CXCR2 inhibitor SB225002 reduces GFAP and CXCR2 in human astrocytes, as shown by immunoblotting (C) and immunofluorescence staining (D).

(E) The CXCR2 inhibitor SB225002 reduces expression of CXCR2 and pNFκB (pP65) in IBC cells.

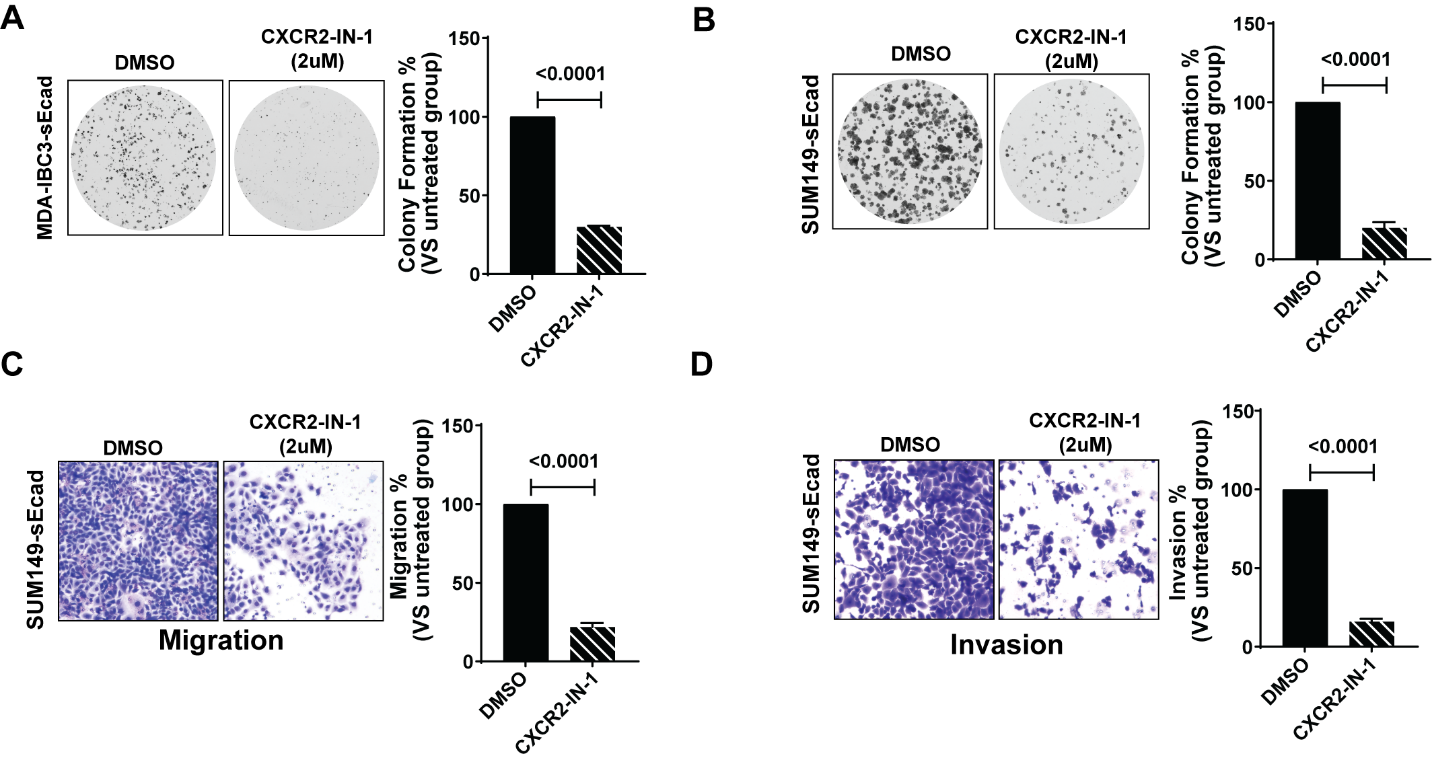

**Figure S10. The inhibitory effect of CXCR2-IN-1 is sEcad-overexpressing IBC cells.**

(A, B) CXCR2-IN-1 inhibits colony formation in sEcad-overexpressing MDA-IBC3 (A) and SUM149 (B) cells. Data are as mean ± SEM of at least three independent experiments.

(C, D) CXCR2-IN-1 inhibits migration (C) and invasion (D) of sEcad-overexpressing SUM149 cells. Data are mean ± SEM of at least three independent experiments.

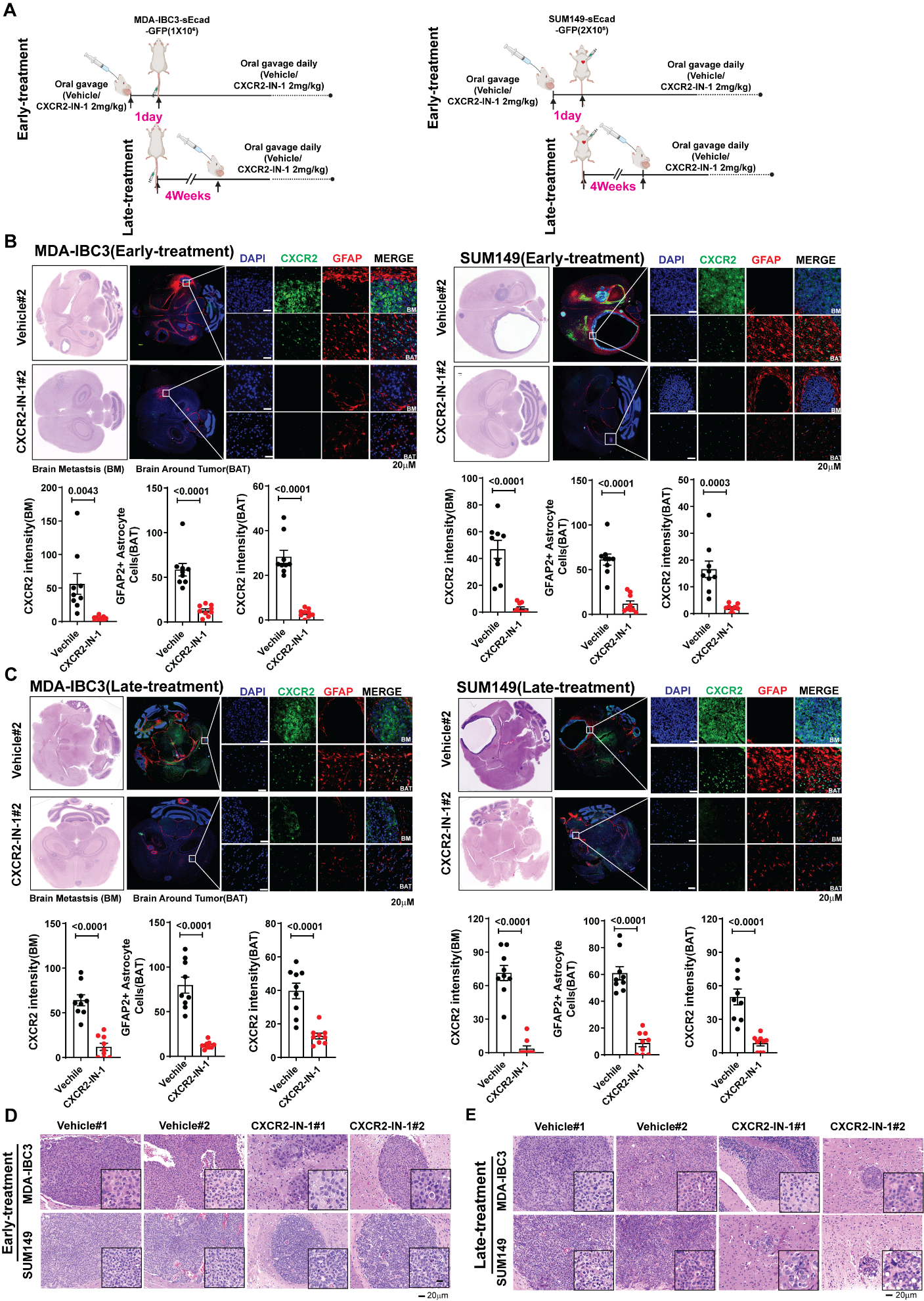

**Figure S11 [previous page]. CXCR2-IN-1reduces metastatic burden and improves survival in IBC models of brain metastasis**.

(A) Schema of the experimental design to treat brain metastasis in two IBC mouse models using two approaches: 'Early Treatment' and 'Late Treatment.'

(B, C) Hematoxylin and eosin and immunofluorescence stains of brain metastasis lesions in mice treated with CXCR2-IN-1 the 'Early treatment' group (B) and the 'Late treatment' group (C) show that CXCR2-IN-1 decreased reactive astrocytes (GFAP+) and CXCR2 expression in both MDA-IBC3-sEcad and SUM149-sEcad brain metastases. Quantitation of CXCR2, GFAP-positive astrocytes. Data are mean ± SEM, n = 3 mice and 3 fields per mouse, t test.

(D, E) Representative hematoxylin and eosin-stained images of brain metastasis lesions generated from MDA-IBC3 and SUM149 tumors in mice treated with vehicle or CXCR2-IN-1, including the 'Early treatment' group (D) and the 'Late treatment' group (E).
