## Supplementary material for "Soluble E-cadherin Drives Brain Metastasis in Inflammatory Breast Cancer": Table1-2

**Table 1. Patient characteristics**

| Covariate | Value or No. (%) |
| --- | --- |
| Patient age, years (n=304), mean±SD | 51.33±11.54 |
| sEcad, ng/mL (n=301), median (IQR) | 65.78 (52.76–80.62) |
| Race |  |
| Black | 28 (9.2%) |
| Other | 8 (2.6%) |
| Hispanic | 30 (9.9%) |
| White | 238 (78.3%) |
| Sex |  |
| Female | 304 (100%) |
| Postmenopausal |  |
| No | 135 (46.1%) |
| Yes | 158 (53.9%) |
| Unknown | 11 |
| Clinical Disease Stage ?at diagnosis |  |
| IIIB | 115 (37.8%) |
| IIIC | 85 (28%) |
| IV | 104 (34.2%) |
| Pathological Disease Stage |  |
| 0-1 | 63 (27.2%) |
| 2 | 27 (11.6%) |
| 3 | 92 (39.7%) |
| 4 | 50 (21.6%) |
| Unknown | 72 |
| ER status |  |
| Negative | 138 (46.3%) |
| Positive | 160 (53.7%) |
| Unknown | 6 |
| PR status |  |
| Negative | 188 (63.5%) |
| Positive | 108 (36.5%) |
| Unknown | 8 |
| HR status (either ER or PR+) |  |
| Negative | 132(44.4%) |
| Positive | 165(55.6%) |
| Unknown | 7 |
| HER2 status |  |
| Negative | 179 (61.7%) |
| Positive | 111 (38.3%) |
| Unknown | 14 |
| HR/HER2 status |  |
| HR+/HER2- | 111 (38.5%) |
| HR+/HER2+ | 50 (17.4%) |
| HR-/HER2+ | 59 (20.5%) |
| HR-/HER2- | 68 (23.6%) |
| Unknown | 16 |
| Histological Grade |  |
| I | 3 (1.1%) |
| II | 62 (22%) |
| III | 217 (77%) |
| Unknown | 22 |
| Lymphatic Invasion |  |
| No | 106 (42.7%) |
| Yes | 142 (57.3%) |
| Unknown | 56 |
| Vascular Invasion |  |
| No | 108 (43.5%) |
| Yes | 140 (56.5%) |
| Unknown | 56 |
| Response to Neoadjuvant Chemo |  |
| ADJ | 1 (0.7%) |
| Complete clinical response | 4 (2.9%) |
| Complete response | 24 (17.1%) |
| Minimal response | 17 (12.1%) |
| Progressive disease | 7 (5%) |
| Partial response | 78 (55.7%) |
| Stable disease | 9 (6.4%) |
| N/A | 164 |
| Neoadj_And_Response |  |
| No neo or no CR | 225 (90.4%) |
| CR | 24 (9.6%) |
| Neo but unknown response | 55 |
| Adjuvant Chemo |  |
| No | 253 (83.2%) |
| Yes | 51 (16.8%) |
| Neoadjuvant Radiation |  |
| No | 300 (98.7%) |
| Yes | 4 (1.3%) |
| Adjuvant Radiation |  |
| No | 141 (46.4%) |
| Yes | 163 (53.6%) |
| sEcad* |  |
| ≤95 | 260 (86.4%) |
| >95 | 41 (13.6%) |
| Missing | 3 |

*99.8 ng/mL was the 90th percentile for sEcad levels.

Abbreviations: sEcad, soluble E-cadherin; ER, estrogen receptor; PR, progesterone receptor; Chemo, chemotherapy; ADJ, adjuvant; N/A, not applicable; CR, complete response; Neo, neoadjuvant (pre-operative) chemotherapy.

**Tables 2+3. Univariate Cox regression analysis for overall and breast-cancer specific deaths**

| Overall Survival* BC-Specific Survival**  BC-Specific Survival (120 BC deaths) | | | | | |
| --- | --- | --- | --- | --- | --- |
| **Covariate** | **HR (95% CI)** | ***P* Value** |  | **HR (95% CI)** | ***P* Value** |
| Age |  |  |  |  |  |
| ≤50 years | 1.000 |  |  | 1.000 |  |
| >50 years | 0.997 (0.703-1.415) | 0.9883 |  | 0.895 (0.625-1.281) | 0.5441 |
| Race |  |  |  |  |  |
| Not Black | 1.000 |  |  | 1.000 |  |
| Black | 1.793 (1.075-2.991) | 0.0253 |  | 1.921 (1.149-3.212) | 0.0128 |
| Postmenopausal |  |  |  |  |  |
| No | 1.000 |  |  | 1.000 |  |
| Yes | 1.021 (0.718-1.453) | 0.9068 |  | 0.980 (0.683-1.408) | 0.9149 |
| Clinical Disease Stage |  |  |  |  |  |
| IIIB | 1.000 |  |  | 1.000 |  |
| IIIC | 1.947 (1.225-3.092) | 0.0048 |  | 1.940 (1.193-3.156) | 0.0076 |
| IV | 2.406 (1.552-3.730) | <.0001 |  | 2.661 (1.693-4.184) | <0.0001 |
| Pathological Stage |  |  |  |  |  |
| 0-1 | 1.000 |  |  | 1.000 |  |
| 2 | 1.976 (0.818-4.773) | 0.1300 |  | 2.145 (0.827-5.563) | 0.1167 |
| 3 | 3.097 (1.590-6.031) | 0.0009 |  | 3.430 (1.654-7.116) | 0.0009 |
| 4 | 2.769 (1.350-5.682) | 0.0055 |  | 3.386 (1.566-7.319) | 0.0019 |
| Estrogen Receptor Status |  |  |  |  |  |
| Negative | 1.000 |  |  | 1.000 |  |
| Positive | 0.673 (0.473-0.958) | 0.0280 |  | 0.656 (0.456-0.943) | 0.0230 |
| Progesterone Receptor Status |  |  |  |  |  |
| Negative | 1.000 |  |  | 1.000 |  |
| Positive | 0.456 (0.302-0.689) | 0.0002 |  | 0.471 (0.309-0.718) | 0.0005 |
| Hormone Receptor Status |  |  |  |  |  |
| Negative | 1.000 |  |  | 1.000 |  |
| Positive | 0.683 (0.480-0.973) | 0.0348 |  | 0.669 (0.465-0.963) | 0.0305 |
| HER2 Status |  |  |  |  |  |
| Negative | 1.000 |  |  | 1.000 |  |
| Positive | 0.277 (0.174-0.441) | <0.0001 |  | 0.281 (0.174-0.453) | <0.0001 |
| HR/HER2 Status |  |  |  |  |  |
| HR+/HER2– | 1.000 |  |  | 1.000 |  |
| HR+/HER2+ | 0.392 (0.198-0.775) | 0.0071 |  | 0.376 (0.183-0.770) | 0.0075 |
| HR–/HER2+ | 0.383 (0.203-0.723) | 0.0031 |  | 0.407 (0.214-0.772) | 0.0059 |
| HR–/HER2– | 2.137 (1.427-3.200) | 0.0002 |  | 2.148 (1.416-3.260) | 0.0003 |
| Grade |  |  |  |  |  |
| I-II | 1.000 |  |  | 1.000 |  |
| III | 1.089 (0.700-1.695) | 0.7058 |  | 1.124 (0.709-1.780) | 0.6194 |
| Lymphatic Invasion |  |  |  |  |  |
| Negative | 1.000 |  |  | 1.000 |  |
| Positive | 1.945 (1.266-2.989) | 0.0024 |  | 2.152 (1.367-3.388) | 0.0009 |
| Vascular Invasion |  |  |  |  |  |
| Negative | 1.000 |  |  | 1.000 |  |
| Positive | 2.027 (1.319-3.112) | 0.0013 |  | 2.241 (1.424-3.528) | 0.0005 |
| Response to Neoadjuvant Chemo |  |  |  |  |  |
| No | 1.000 |  |  | 1.000 |  |
| Res | 0.190 (0.060-0.599) | 0.0046 |  | 0.131 (0.032-0.532) | 0.0045 |
| Receipt of Adjuvant Chemo |  |  |  |  |  |
| No | 1.000 |  |  | 1.000 |  |
| Yes | 0.451 (0.262-0.779) | 0.0043 |  | 0.442 (0.251-0.776) | 0.0045 |
| Receipt of Neoadjuvant Radiation |  |  |  |  |  |
| No | 1.000 |  |  | 1.000 |  |
| Yes | 1.770 (0.561-5.587) | 0.3305 |  | 1.855 (0.587-5.864) | 0.2925 |
| Receipt of Adjuvant Radiation |  |  |  |  |  |
| No | 1.000 |  |  | 1.000 |  |
| Yes | 0.368 (0.257-0.525) | <0.0001 |  | 0.339 (0.234-0.491) | <0.0001 |
| sEcad |  |  |  |  |  |
| ≤95 | 1.000 |  |  | 1.000 |  |
| >95 | 1.596 (1.007-2.529) | 0.0466 |  | 1.614 (1.007-2.587) | 0.0466 |

*127 deaths

**120 deaths

Abbreviations: HR, hazard ratio; CI, confidence interval; Chemo, chemotherapy sEcad, soluble E-cadherin.
